# An Expanded Repertoire of tRNA Sources for Cell-Free Protein Synthesis

**DOI:** 10.1101/2025.08.20.671396

**Authors:** Evan M. Kalb, Russel M. Vincent, Aaron E. Engelhart, George M. Church, Katarzyna P. Adamala

**Affiliations:** Department of Genetics, Cell Biology and Development, University of Minnesota, Minneapolis, MN US; Department of Genetics, Harvard Medical School, Boston, MA 02115, USA; Wyss Institute for Biologically Inspired Engineering, Harvard University, Boston, MA 02215, USA; Macquarie Medical School, Faculty of Medicine, Health & Human Sciences, Macquarie University, Sydney, NSW 2109, Australia

## Abstract

Cell-free expression systems (CFE) are flexible protein translation platforms that simplify the central dogma into an accessible reaction space. Within these systems, bulk transfer RNAs (tRNAs) are critical substrates which deliver amino acids to the elongating ribosome. For years, CFE systems were completed with commercially available tRNA isolated from *E. coli* MRE600. All commercial sources of tRNA have since been discontinued, jeopardizing future work in all applications of cell-free translation.

Here, we address this need by repurposing previously described tRNA isolation methods to produce tRNAs suitable for CFE applications. We isolated the tRNA pools of *E. coli* strains A19, BL21(DE3), and Rosetta2 BL21(DE3), finding A19 tRNAs but not BL21(DE3) or Rosetta2 BL21(DE3) capable of robust *in vitro* translation. We determined the abundances of individual tRNAs using tRNA-seq, finding BL21(DE3) and Rosetta2 BL21(DE3) contained outsized abundances of several tRNAs, compromising translation activity. Using codon optimization strategies which align codon usage to tRNA abundance, we were able to mitigate the impact of misaligned tRNA abundances. We extended these studies to *V. natriegens*, a promising platform for synthetic biology and CFE. We find that neither exogenous *V. natriegens* tRNAs nor codon optimization are viable options to improve translation yields. Our work here highlights the importance of tRNA abundance within the context of CFE, and simultaneously addresses a critical challenge within cell-free translation.

## Introduction

Cell-free protein expression systems (CFE)^1^ reduce the complexities found within a living central dogma by separating gene expression from the burden of organismal homeostasis. Without the constraints of living systems, CFEs present an open biochemical platform that allows the direct manipulation of the gene expression. Such control has enabled the study of protein translation that would otherwise be difficult in living systems, including the elucidation of the universal genetic code^2–4^, introduction of non-canonical amino acids^5–9^, and the assembly and prototyping of complex metabolic pathways^10–13^.

CFE systems are broadly classified into two categories. Lysate-based systems^14^ (hereon S30 CFE) utilize the cytosolic components of once-living organisms for convenient and inexpensive protein synthesis. In contrast, Protein synthesis Using Recombinant Elements (PURE)^15^ represents a minimalist model of gene expression by reducing transcription and translation to their core molecular components. In PURE, 36 proteins, ribosomes, tRNAs, and small molecules such as amino acids, NTPs, and phosphate donors are individually synthesized, purified, and reassembled into a defined biochemical reaction. This chemically defined environment provides a foundational platform of transcription and translation, where complexity can be added to its core structure.

Both PURE and CFE based systems have enabled technologies ranging from biosensors^16,17^, on-demand biomanufacturing^18,19^, and therapeutics^20^. Beyond these applications, *in vitro* translation systems represent the bottom-up and middle-out strategies of synthetic cell development, respectively^21^. PURE’s example as a minimalist translation system makes it an ideal reference model for the development of a self-replicating central dogma^22^ and synthetic cells.

The traditional tRNA complement for PURE CFE was the once commercially available *E. coli* MRE600 tRNA isolate^15^. Commercial MRE600 tRNAs are also used to supplement S30 CFE systems^23–26^, including those which employ non-*E. coli* lysates^27–31^. MRE600 tRNAs are no longer commercially available, and the only other commercial alternative, tRNA from *E. coli* W strain, exhibits comparatively poor translation activity^32^. Recently, *E. coli* W strain tRNAs have also become unavailable. Without access to convenient tRNA sources, PURE CFE development is fundamentally compromised, representing a dire threat to the CFE community.

We address this critical bottleneck by examining *E. coli* strains A19, BL21(DE3), and Rosetta2 BL21(DE3) as tRNA sources to complement CFEs. We first evaluated the translational capabilities of the candidate tRNA pools in PURE CFE by translating sfGFP and NanoLuc. We found A19 tRNAs capable of equal translation to commercial MRE600 tRNAs, while BL21(DE3), and Rosetta2 BL21(DE3) activities lagged.

We determined translation yields in S30 CFE systems assembled from either A19 or Rosetta2 BL21(DE3) strain lysates, finding them identical unless codon bias of the sfGFP reporter was significantly altered. Seeking to explain the differing activities, we used tRNA-seq to determine tRNA abundances of each strain, finding BL21(DE3), and Rosetta2 BL21(DE3) contained disproportionately high abundances of tRNA^Pro^_GGG_, tRNA^eMet^, and tRNA^Leu^ compared to A19. With knowledge of strain tRNA abundances, we designed a simple codon optimization strategy which reduces the impact of underutilized tRNAs. Our approach, called tRNA-inclusive codon optimization or TICO, generated constructs with improved yield in PURE CFE.

Finally, we utilized isolated *V. natriegens* tRNAs in both PURE and S30 CFE reactions to examine the potential for exogenous tRNAs to improve *V. natriegens* S30 CFE systems. We found that the *V. natriegens* tRNAs are poor substrates for *E. coli* PURE CFE despite their shared sequence similarity. In contrast, S30 CFE composed of *V. natriegens* lysates preferred *E. coli* optimized constructs. Moreover, supplemental *V. natriegens* tRNA was largely ineffective in improving the yields within the *V. natriegens* S30 reactions.

Our results show that strain-specific tRNA preferences determine translation efficiency in PURE and S30 CFE systems. We not only address reagent scarcity for PURE and S30 CFE reactions, but also provide a conceptual framework for future codon optimization strategies within cell-free contexts. These findings place tRNA pool composition and codon usage as central levers for systematically enhancing translation efficiency and yield in PURE CFE.

## Results

### Strain selection for candidate *E. coli* tRNA sources

CFE reactions typically use *E. coli* MRE600 tRNA due its once widespread commercial availability. *E. coli* MRE600 is deficient in RNase I, allowing the accumulation of RNA products^33^. Because of this, MRE600 is the *de facto* strain for the isolation of both ribosomes and tRNAs for *in vitro* studies^34^. Still, there are significant challenges in utilizing MRE600 as a source for tRNAs for CFEs. MRE600 is a *Shigella*-like strain of *E. coli*^34^, and is designated by some entities as Biosafety Level II, including the BacDive database (https://bacdive.dsmz.de/strain/4468). This classification poses a challenge for labs unable to accommodate these organisms.

We considered alternative *E. coli* strains as potential tRNA sources for PURE and S30 CFE. One alternative to MRE600 is *E. coli* A19, a K12 derivative that is also RNase I deficient^33^. Because of this, A19 is a staple source of ribosomes for homemade PURE formulations^35^. A19 also contains mutations in *relA* and *spoT* which are key initiators of the stringent response, potentially making it resistant to wholesale degradation of stable RNAs under stressed conditions. Notably, extracts generated from stressed A19 cells are equally as active those harvested at the typical mid-log phase^36^. Additionally, A19 is a popular lysate for S30 CFE^36–39^. We also evaluated BL21 (DE3), a strain commonly used for protein expression and whose lysates are most often used to generate lysates for S30 CFE^25^. We were similarly interested in Rosetta2, a derivative of BL21 (DE3) (hereon Rosetta) which is used to improve expression of eukaryotic proteins^40^. Rosetta contains the pRARE2 plasmid which includes the genes required to transcribe otherwise scarce *E. coli* tRNAs. This feature has made Rosetta another common lysate for *E. coli* S30 CFE systems^41,42^.

### Strain choice determines the translational potential of tRNA pools

We isolated tRNAs from *E. coli* MRE600, A19, BL21, and Rosetta strains using a protocol described by Avcilar-Kucukgoze et. al^43^ which, in principle, shares the isolation approaches of Zubay^44^, Von Ehrenstein^45^, and Cayama et al.^46^. Since tRNA abundance can influence translation outcomes and may change in response to environmental conditions such as starvation^47^, we harvested tRNAs at the early to mid-log phase typical of lysate preparations for S30 CFE^48^. Generally, all strain sources produced high yields of tRNA isolates ranging from 5-20 mg. Notably, Rosetta produced greater yields than either A19 or BL21, perhaps due to the expression of the pRARE2 tRNAs. We first determined tRNA quality using denaturing urea-PAGE (**Supplemental Information S1)**. Recovered tRNAs were pure and free of DNA and 16S and 23S rRNA contaminants. Along with tRNAs, transfer-messenger RNA (tmRNA) and 5S rRNA were observed, and both were also present in commercial MRE600 tRNAs **(Supplemental Information S1)**.

We determined the activity of the recovered tRNA pools by using a commercially available PURE CFE system which separates tRNAs from the energy mix, allowing users discrete control of the tRNA pool. We completed these reactions with tRNA pools isolated from either MRE600, A19, Rosetta, or BL21 and tasked the pools with the translation of pJL1 *E. coli* sfGFP. Both kit-supplied tRNAs and the typical commercial MRE600 tRNAs from Roche served as positive controls. sfGFP yields from reactions utilizing A19 (0.256 ± 0.037 mg/mL) and BL21 (0.270 ± 0.005 mg/mL) tRNA pools were essentially equal to both the kit-supplied tRNA from New England Biolabs (NEB, 0.269 ± 0.035 mg/mL) and commercial Roche MRE600 tRNA (0.245 ± 0.007 mg/mL), showing no statistically significant difference in yields **(Figure 1a)**. In contrast, Rosetta tRNA achieved translation yields ∼80% (0.214 ± 0.015 mg/mL) of NEB tRNAs, suggesting that supplemental rare, plasmid-borne tRNAs might be inhibitory to translation. Surprisingly, in-house prepared MRE600 tRNAs failed to produce any observable sfGFP and further attempts to recover activity through urea-PAGE gel-based purification and dialysis also failed **(data not shown)**. Consequently, we did not pursue in-house MRE600 tRNAs further.

**Figure 1.**
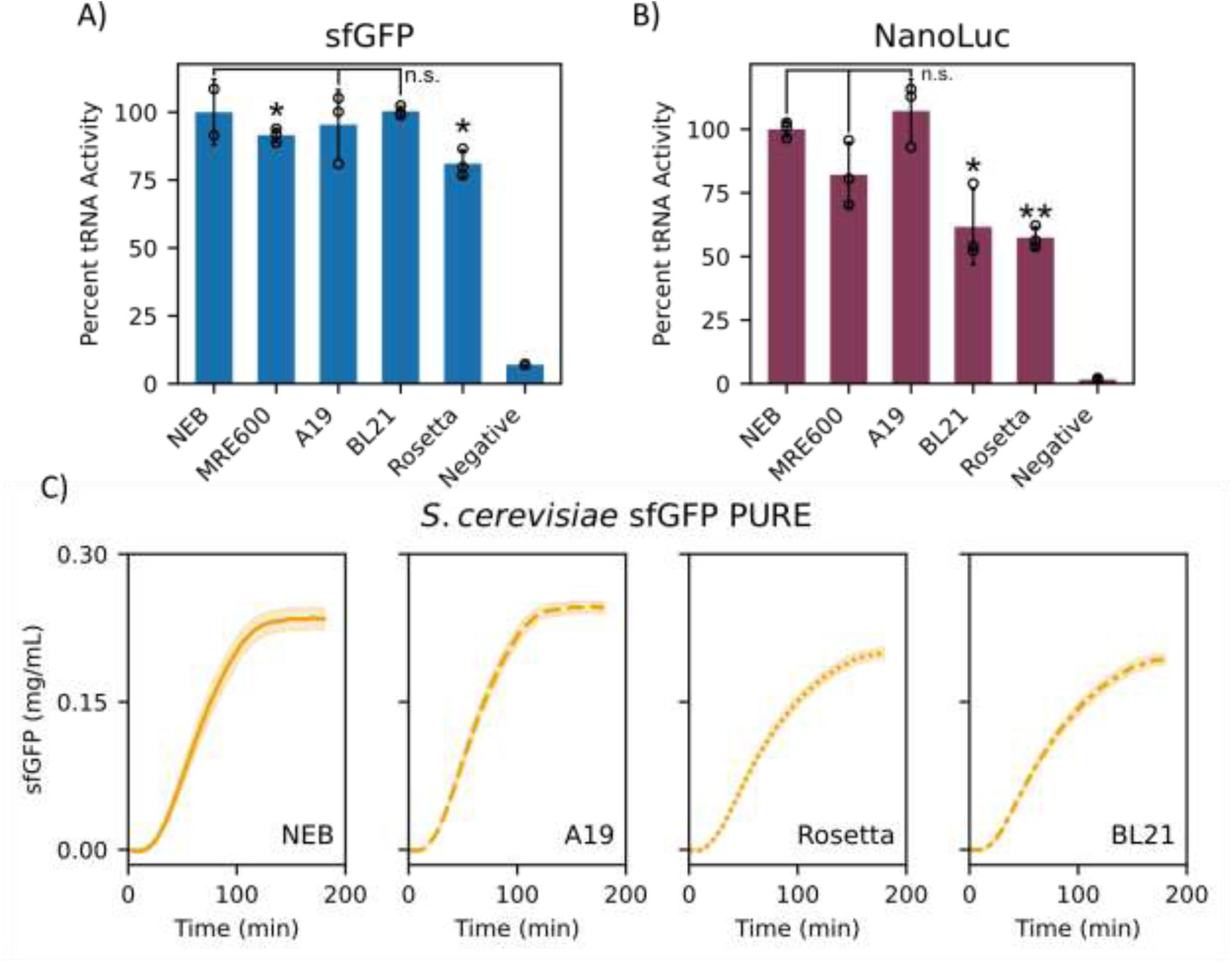
Isolated *E. coli* tRNA pools complete ΔtRNA PURE CFE. Isolated *E. coli* tRNAs facilitate translation in ΔtRNA PURE. **A)** ΔtRNA PURE CFE translating pJL1 *E. coli* sfGFP completed with tRNA pools harvested from either A19, BL21, or Rosetta *E. coli* strains. Commercial MRE600 and kit-supplied NEB tRNAs served as positive controls while no tRNA added served as a negative control. Reactions yields were normalized to end-point relative fluorescence units of NEB tRNA completed PURE reactions. A19 and BL21 tRNA pools exhibit no difference in translation capabilities compared to NEB tRNA. Commercial MRE600 and Rosetta tRNAs demonstrate modest, but statistically significant reduction in translation performance (p-values of 0.029 and 0.022, respectively). Open circles represent individual replicates and error bars represent standard deviation of three replicates. Significance was calculated from a one-sample t-test compared to NEB. **B)** Same as in A except translating *Renilla reniformis* NanoLuc. Translation yields were determined from relative luminescence values tracked for 10 minutes after mixing with NanoGlo Extracellular reagent (Promega, Madison, Wisconsin). Yields were normalized to NEB tRNA end-point luminescence. BL21 and Rosetta pools demonstrate significant losses in translation compared to kit-provided tRNAs (p-values of 0.046 and 0.0034, respectively. **C)** Same as A except for a time-course translation of ΔtRNA PURE CFE translating *S. cerevisiae* optimized sfGFP. Shaded regions represent standard deviation of mean of three replicates.

We extended PURE CFE reactions to the translation of *Renilla reniformis* NanoLuc. Again, A19 tRNAs maintained the same translation capabilities of both the kit-supplied and commercial Roche MRE600 tRNAs as determined by end-point luminescence assays **(Figure 1b)**. However, both BL21 and Rosetta tRNA pools suffered diminished yields relative to kit-provided tRNAs (62% and 57%, p-values of 0.046 and 0.003, respectively). Since BL21 and Rosetta share the same parent strain, we suspected these differences in translation capabilities might stem from the tRNA abundances produced by the BL21 genome.

Given that Rosetta is often used to translate eukaryotic proteins, we wondered if its tRNA pool could outperform either BL21 or A19 tRNA pools if the coding sequence of sfGFP aligned to a eukaryotic codon bias. Using a commercial gene synthesis service’s in-house optimization algorithm, we generated an sfGFP construct codon optimized for expression in *Saccharomyces cerevisiae* while maintaining the non-coding regulatory elements such as 5’-UTR, ribosome binding site, and 3’-UTR. The resulting construct increased the codon frequencies of ten codons that would benefit from supplemental tRNAs provided by the pRARE2 plasmid **(Supplemental Information S2)**. To our surprise, Rosetta tRNAs failed to significantly improve translation compared to BL21, and both BL21 and Rosetta tRNA pools lagged behind A19 and kit tRNAs **(Figure 1c)**. BL21 and Rosetta tRNA pools achieved yields relative to NEB tRNAs at 82.6% ± 4.4% (0.194 ± 0.006 mg/mL) and 85.2% ± 4.4% (0.200 ± 0.005 mg/mL), respectively, while A19 effectively matched NEB tRNA yields (A19: 0.246 ± 0.006 mg/mL; NEB: 0.235 ± 0.010 mg/mL). These results suggest that both BL21 and Rosetta tRNA pools are inadequate to support efficient translation in PURE CFE under our conditions. Moreover, rare tRNAs provided by pRARE2 plasmid are incapable of addressing these inefficiencies, even when translation is biased towards eukaryotic codon preferences. In contrast, A19 and kit-supplied tRNAs are equally capable in addressing the translational demands of *S. cerevisiae* sfGFP.

### Exogenous tRNAs variably improve lysate-based CFE

Despite the retention of cytosolic tRNAs, commercial MRE600 tRNAs are typically used to supplement lysate-based CFE systems^25,48^. Previous work suggests exogenous tRNAs may not be necessary for high translation yields^49^, and it is unclear under what circumstances exogenous tRNAs are beneficial. Given the disparate translation capabilities of A19 and Rosetta tRNA pools in PURE, we wondered if similar trends would be reflected in S30 CFE reactions, especially when translating *S. cerevisiae* sfGFP. Should deficiencies arise, we were curious to see if exogenous tRNAs would enable greater translation yields. Thus, we generated lysates from both A19 and Rosetta *E. coli* and prepared S30 CFE reactions to translate either *E. coli* or *S. cerevisiae* sfGFP. Additionally, a supplement of 0.25 mg/mL of either A19 or Rosetta tRNAs were added to evaluate the impact of exogenous tRNA in translation.

Both A19 and Rosetta S30 CFE reactions lacking supplemental tRNAs translated *E. coli* optimized pJL1 sfGFP equally well, indicating that both endogenous tRNA pools were capable of efficient expression **(Figure 2a-b).** Additionally, there were only minor improvements in *E. coli* sfGFP translation yields in both lysates when either A19 or Rosetta tRNA pools were added to the system **(Figure 2a-b).** However, in *S. cerevisiae* sfGFP translation, there was a far greater difference in expression capabilities between the lysates. Rosetta S30 CFE was able to translate *S. cerevisiae* sfGFP at nearly ∼80% of its *E. coli* sfGFP yield, and additional A19 tRNAs narrowly improved yields **(Figure 2b)**. In contrast, A19 S30 struggled to translate *S. cerevisiae* sfGFP, generating roughly one third of the sfGFP encoded by the *E. coli* optimized codons within the same lysate **(Figure 2a)**. A19 S30 supplemented with A19 tRNAs marginally recovered *S. cerevisiae* sfGFP expression (12.1% ± 4.4%), but supplemental Rosetta tRNAs improved the yield appreciably (35.3% ± 2.6%) **(Figure 2a)**.

**Figure 2.**
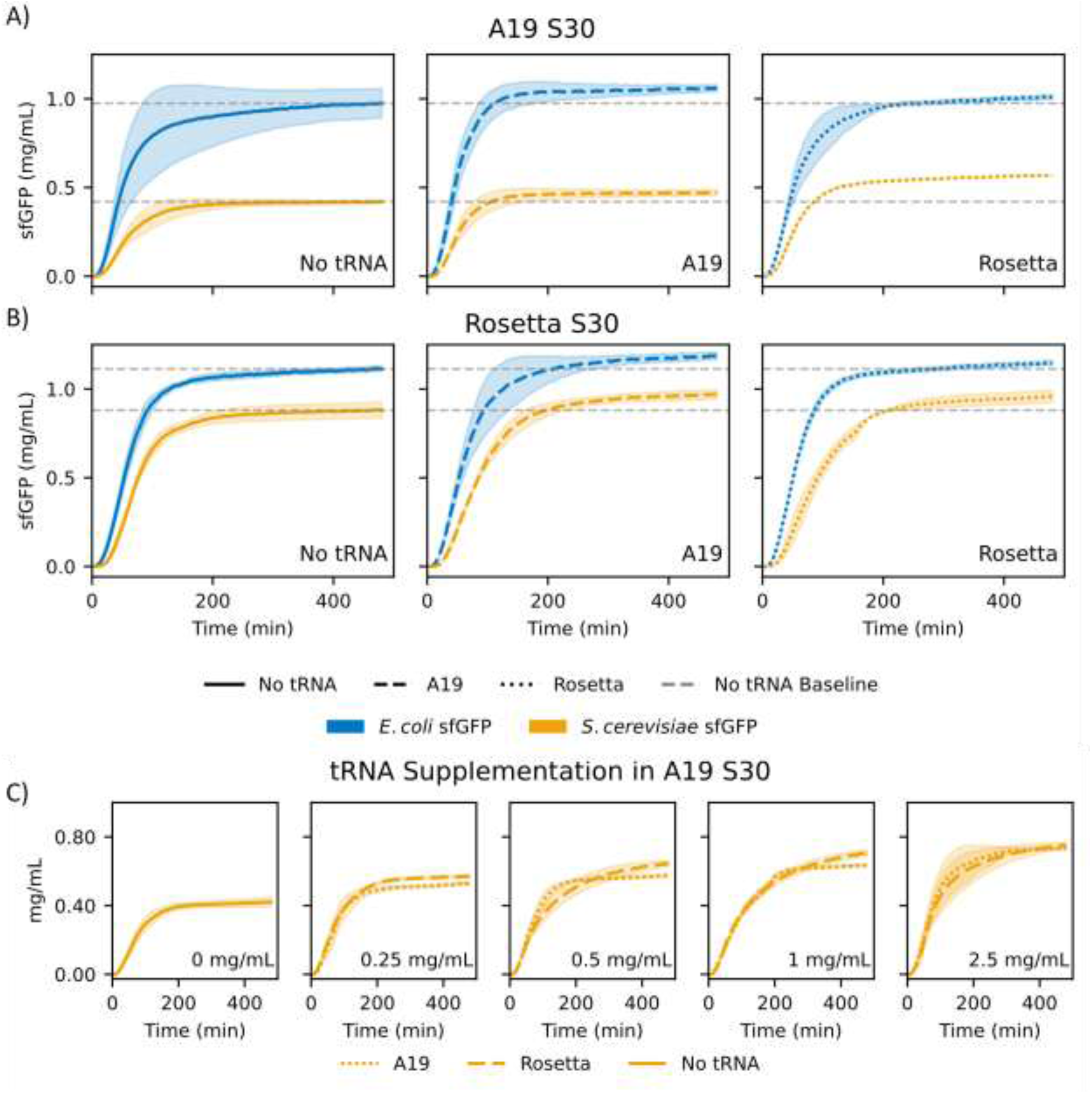
Supplementation of exogenous tRNA pools selectively improves S30 CFE. **A)** A19 S30 CFE translating pJL1 *E. coli* sfGFP (blue) or pJL1 codon optimized to *S. cerevisiae* sfGFP (gold) with the supplementation of either A19 or Rosetta tRNAs at a final concentration of 0.2 mg/mL. Dashed grey lines represent the end-point yield baseline for a control containing no supplemental tRNA. **B)** Same as in A, except for the CFE utilizing a Rosetta S30 lysate. **C)** A19 S30 translating codon optimized to *S. cerevisiae* sfGFP (gold) with increasing final concentrations of either supplemental A19 (dotted lines) or Rosetta tRNA (dashed lines). RFU values were converted into mg/mL protein produced using a standard curve **(see Experimental Section)**. Shaded regions represent the standard deviation of mean across three experimental replicates.

These data reveal that endogenous tRNAs in A19 S30 cannot sustain the demands of *S. cerevisiae* sfGFP translation. This translational bottleneck is partially alleviated by the exogenous Rosetta tRNA pool but only slightly by the A19 tRNA pool, indicating that the Rosetta pool is indeed supplying rarer tRNAs needed for *S. cerevisiae* sfGFP translation. Although PURE CFE using Rosetta tRNAs achieved reduced yields compared to A19 tRNAs, Rosetta S30 CFE exhibited strong translation efficiency for both *S. cerevisiae* and *E. coli* coding sequences. These results suggest that cytosolic translation components in S30 reactions can better tolerate mismatches in tRNA pool and coding sequences, so long as tRNAs are sufficiently supplied.

Since exogenous Rosetta tRNA aided A19 S30 in *S. cerevisiae* sfGFP translation, we hypothesized that additional exogenous tRNAs may further improve translation yields. Specifically, we supposed that additional A19 or Rosetta tRNAs would both improve the translation of *S. cerevisiae* sfGFP with Rosetta tRNA more efficiently recovering translation due to its expected greater enrichment in rare tRNAs. To test this, we added increasing amounts of exogenous tRNAs sourced from either A19 or Rosetta tRNA to A19 S30 reactions translating *S. cerevisiae* sfGFP. As expected, the end-point yields of *S. cerevisiae* translation improved with each addition of exogenous tRNA, tapering off at final concentration of 2.5 mg/mL supplied tRNAs **(Figure 2c)**. In these reactions, Rosetta tRNAs narrowly outpaced A19 tRNAs, suggesting the A19 pool was still supplying the rarer tRNAs but with a more modest impact. While the maximum of added tRNAs nearly doubled translation yields of unsupplemented A19 S30 CFE (total improvement of 80.2% ± 5.7%), this supplemented reaction still only achieved 80% of the translation yield of the highest yielding Rosetta S30 CFE reaction. These results suggest that supplementation of exogenous tRNAs to lysates improves translation only up to a point, beyond which diminishing returns limit further gains. Thus, endogenous tRNAs cannot be completely superseded with exogenous tRNAs.

### tRNA-seq reveals differences in tRNA abundances within and across species

Given the differences in translation capabilities, we were interested in determining the abundances of individual tRNAs within A19, BL21, Rosetta, and commercial MRE600 tRNA pools. We were also interested in determining the abundances within the tRNA pool of *Vibrio natriegens*, a halophilic, Gram-negative gammaproteobacteria. Because of its rapid doubling time and ease of genetic manipulation^50^, *V. natriegens* has attracted great interest as an alternative chassis for synthetic biology efforts^51^ including heterologous protein expression^52^ and metabolic engineering^51,53^. Despite this potential, *V. natriegens* S30 CFE lysates lag behind *E. coli* S30 CFE translation yields^31,54^, and we wondered if a deeper understanding of its tRNA pool might improve its use in lysate-based CFE. To do so, we isolated tRNAs from *V. natriegens* tRNAs using the same method for *E. coli* tRNA, but were met with comparatively poor yields. Slight modifications to the protocol were needed to recover tRNA in yields sufficient for downstream applications (see **Experimental Section**).

We employed tRNA-seq on isolated tRNAs from *E. coli* A19, BL21, Rosetta, commercial MRE600, and *V. natriegens*, determining both abundance of individual tRNAs in reads per million (RPM) **(Supplemental Information SF3)** and calculated the percent relative abundance of each tRNA within its tRNA pool **(Figure 3a).** tRNA-seq revealed significant differences in relative tRNA abundance across *E. coli* strains. A19 and MRE600 shared similar abundances across most tRNAs except for tRNA^Arg^ which is enriched in MRE600. This supports our observations in PURE CFE where MRE600 and A19 tRNA performed similarly in PURE CFE. Except for a few cases, nearly all tRNAs from A19 and MRE600 exhibited greater abundance (in RPM) than those from BL21 or Rosetta. **(Supplemental Information SF3)**. Instead, both BL21 and Rosetta tRNA pools have dramatically increased abundances of tRNA^eMet^_CAU_, tRNA^Pro^_GGG_, tRNA^Leu^_UAG_, and non-ACG tRNA^Arg^ isoacceptors compared to A19 and MRE600. For example, tRNA^Pro^_GGG_, tRNA^eMet^_CAU_, and tRNA^Leu^_UAG_ occupy 9.8%, 5.6%, and 3.1% of the total BL21 tRNA pool, respectively, compared to 1.5%, 2.0%, and 0.7% in A19 **(Figure 3a)**.

**Figure 3.**
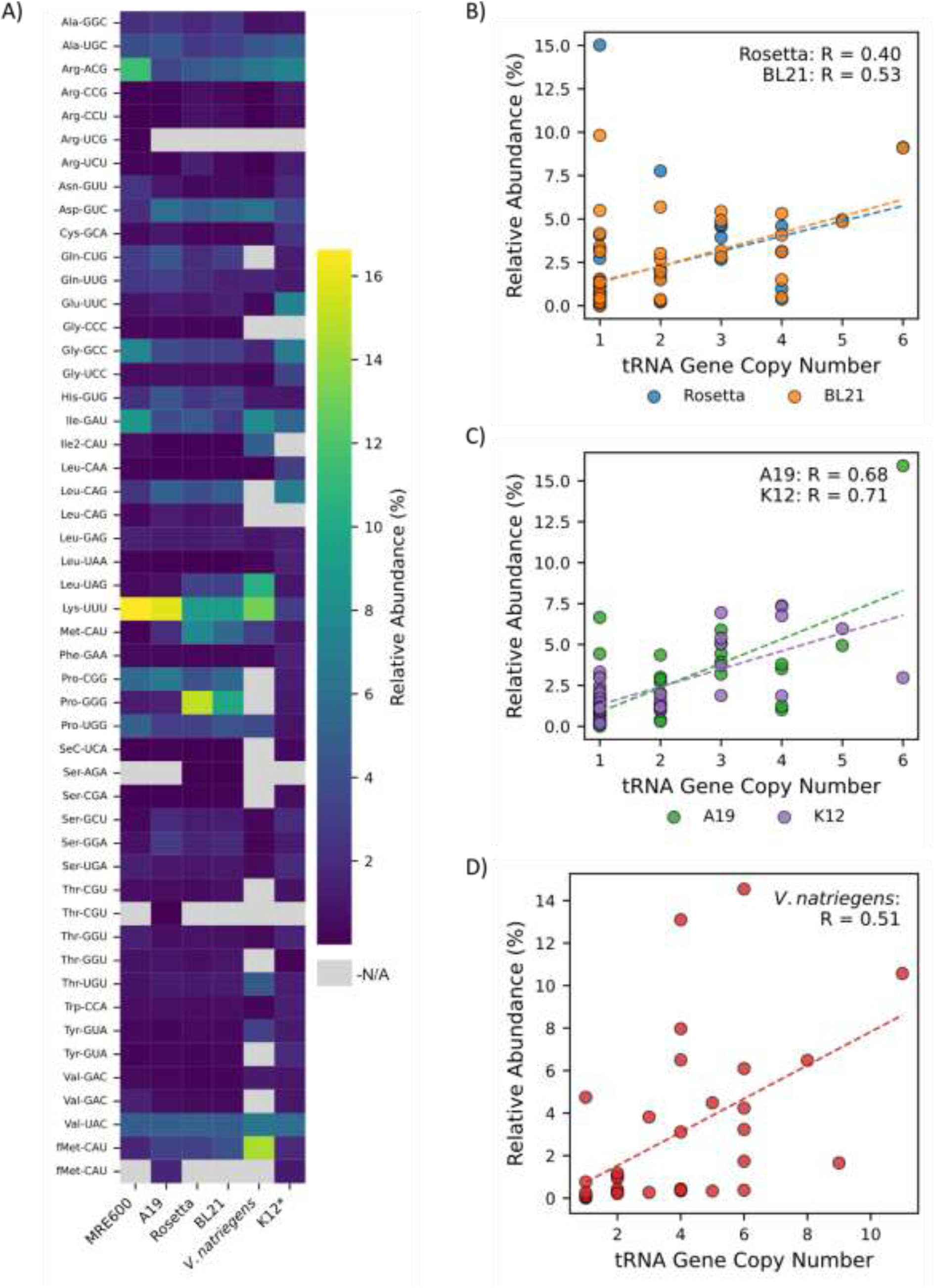
Relative individual tRNA Abundances within *E. coli* and *V. natriegens*. **Relative tRNA abundances differ across *E. coli* strains and *V. natriegens* and poorly correlate to gene copy number. A)** Heatmap of percent relative tRNA abundances for *E. coli* A19, *E. coli* Rosetta(DE3), *E. coli* BL21(DE3), and *V. natriegens* tRNA pools isolated using acid-phenol extraction and sequenced using tRNA-seq. For each strain, percent relative abundances for each unique tRNA were calculated from the mapped reads per million of each unique tRNA divided against the sum reads per million of all tRNAs of the strain. Average reads per million were determined from three biological replicates. For MRE600, tRNA-seq was performed on five experimental replicates of the same commercial source of tRNA from Roche. Genomes lacking tRNA genes according to tRNA-scan were excluded and are represented as grey boxes. **\***For *E. coli* K12, percent relative abundances are taken from Dong et. al 1996^56^; tRNAs undetected in their study are also represented as grey boxes. **B)** Correlation plot of percent relative abundance for unique tRNA reads for Rosetta (blue) and BL21 (orange) strains against their corresponding unique gene copy numbers within their shared BL21 genome. **C)** Correlation plot of percent relative abundance for unique tRNA reads for A19 (green) and K12 (purple) strains against their corresponding unique gene copy numbers within their shared K12 genome. **D)** Correlation plot of percent relative abundance for unique tRNA reads for *V. natriegens* against corresponding unique gene copy numbers within the *V. natriegens* genome. tRNA gene copy numbers were calculated from tRNA-scan predictions^65^. Colored dashed lines correspond to the linear regression. R represents Pearson correlation coefficient.

Sequencing also revealed to what extent pRARE2 increases tRNA abundances within Rosetta. pRARE2 encodes the tRNA genes *metT, argW, argU, argX*, *proL, leuW, IleX, thrT, thrU,* and *glyT* which are expressed under their native promoters. This results in tRNAs which are already highly abundant in BL21 to have an even greater abundance in Rosetta. For example, the pRARE2 plasmid increased percent abundances for tRNA^Pro^_GGG_ from 9.8% to 15.0% and for tRNA^eMet^ from 5.6% to 7.7% **(Fig. 3a)**. The pRARE2 plasmid also increased abundances of rare tRNA^Arg^ isoacceptors in RPM compared to BL21 (**Supplementary Information SF3)**, but these increases were ultimately diluted out to the contributions of tRNA^Pro^_GGG_ and tRNA^eMet^_CAU_. pRARE2 also includes the gene *argN5* (encoding tRNA^Arg^_UCG_) which is absent in the BL21 genome. BLAST identified this as a tRNA found within the genome of the pathogenic *E. coli* O157:H7 strain. Only fragments mapped to *argN5*, suggesting that BL21 is incapable of correctly processing this tRNA.

We suspected the exceedingly high abundances of tRNA^Pro^_GGG_, tRNA^eMet^_CAU_, and tRNA^Leu^_UAG_ transcribed from the BL21 genome (and exacerbated by the pRARE2 plasmid) likely explain the reduced translational yields we observed in PURE CFE. Furthermore, tRNA-seq sheds light on the improved translation of *S. cerevisiae* sfGFP by Rosetta S30 CFE compared to A19. Codon optimization of pJL1 *E. coli* sfGFP to *S. cerevisiae* resulted in an increase of codon frequencies of AGA (+1.66%) and AGG (+0.83%) (**Supplementary Information SF2)**. AGA and AGG codons are decoded by the rare tRNAs, tRNA^Arg^ and tRNA^Arg^ which in the A19 strain occupy only 0.18% and 0.08% of the total tRNA pool, respectively. In contrast, Rosetta strain pools increase tRNA^Arg^ and tRNA^Arg^ abundances by nearly 10-fold to 1.5% and 0.85% of the total tRNA pool respectively, compensating for the increased codon usage. In *S. cerevisiae* sfGFP, leucine codon preferences are more dramatically shifted with an increase in CUA (1.24%), CUU (2.07%), and UUA (1.65%) codons. In Rosetta, tRNA^Leu^_UAG_ is present at 3.26% of the total tRNA pool compared to the 0.65% observed in A19. Notably, in *E. coli*, tRNA^Leu^_UAG_ contains a modified U34 which enables the decoding of all four CUN codons^55^, and its increased abundance within the Rosetta pool can address the increased codon demands of *S. cerevisiae* sfGFP. This is especially true for CUA codons which are exclusively decoded by tRNA^Leu^_UAG_ in *E. coli*.

A19 is derived from *E. coli* K12, the tRNA pool of which has been previously elucidated using oligonucleotide probing by Dong et. al.^56^ This provides another experimental comparison to our tRNA-seq data. A19 tRNA abundances largely agreed with this earlier work **(Figure 3a)**, with a few notable exceptions. Because of sequence similarities, Dong et al. were unable to separate tRNA^IleII^_CAU_ from its sibling isoacceptor, tRNA^IleI^_GAU_, obscuring its abundance. Additionally, some tRNAs were underrepresented in the Dong dataset, such as tRNA^Lys^, tRNA^Pro^ isoacceptors, and tRNA^Gln^, perhaps due to inefficient probe hybridization. Considering the shared tRNA abundance similarities between A19 and *E. coli* K12, we speculate any K12-derived strain could serve as an effective source of tRNAs for PURE CFE.

The tRNA-seq experiments uncovered surprising tRNA preferences in *V. natriegens*, illuminating unique features of its translation strategy. Curiously, *V. natriegens* has an extraordinarily large abundance of initiator tRNA^fMet^_CAU_, occupying nearly 15% of its entire tRNA pool, compared to the 3-4% found among *E. coli* strains **(Figure 3a)**. For elongation tRNAs, *V. natriegens* generally employs a succinct tRNA pool where tRNA abundance is biased towards a single tRNA among its sibling isoacceptors **(Figure 3a)**. For example, tRNA^Leu^_CAG_, the preferred and most highly abundant tRNA for leucine decoding in *E. coli* is absent in *V. natriegens.* Instead, *V. natriegens* employs tRNA^Leu^_UAG_ to decode CUA and CUG codons. This trend is shared across other tRNAs which contain a 5’-CNN-3’ anticodon; *V. natriegens* lacks the genes necessary to transcribe tRNA^Ser^_CGA_, tRNA^Gly^_CCC_, tRNA^Gln^, tRNA^Thr^_CGU_, and tRNA^Pro^_CGG_ found within *E. coli*. Consequently, tRNAs containing 5’-UNN-3’ are generally enriched in *V. natriegens*. For example, tRNA^Leu^_UAG_ and tRNA^Thr^_UGU_ abundances dominate among their respective isoacceptors. Similarly, *V. natriegens* relies on a single tRNA^Pro^_UGG_ to decode all four proline codons. For these codons, expanded decoding is supported by superwobbling, where tRNAs are able to engage with codons beyond the wobble rules proposed by Crick^57^. Specifically, uridines within the wobble position of anticodons (position 34) have been shown to decode all four wobble nucleotides within codons^55,58^. Superwobbling enables a reduced tRNA gene set needed to decode the 64 codons, and organisms containing minimal genomes exploit superwobbling to reduce tRNA requirements^59^.

The relative tRNA abundance data gained from tRNA-seq also allowed us to evaluate assumptions made in codon optimization strategies. Most codon optimization efforts attempt to maximize Codon Adaptation Index (CAI) which estimates gene optimality by comparing a gene’s codon usage relative to highly expressed genes of the host organism^60,61^. CAI has inspired other indices, such as the tRNA Adaptation Index (tAI), which considers optimality within the context of tRNA abundance^62^. In tAI, tRNA abundance is inferred from tRNA gene copy number (tGCN) and has been previously considered to correlate strongly with tRNA abundance^63,64^. We were curious if calculated tRNA abundances from our sequencing efforts would similarly correlate with tGCN.

We generated correlation plots comparing relative tRNA abundances for each unique tRNA against the tGCN annotated using tRNA-scan^65^. In general, tGCN represented a poor approximation of tRNA abundances. Rosetta and BL21 both demonstrated moderate, positive correlation between percent abundances and tGCN (R=0.40 and R=0.53, respectively **(Fig 3b)**. For Rosetta, tRNAs expressed from the pRARE2 plasmid reduced correlation, and Pearson correlation coefficient which was unchanged if pRARE2 copy number is considered **(Supplementary Fig SF4)**. In contrast, A19 and K12 tRNAs pools were more strongly correlated to tGCN, showing Pearson correlation coefficients of 0.68 and 0.71, respectively **(Fig 3c)**.

*V. natriegens* tRNA abundances also had moderate correlation with tGCN (R=0.51, **Fig 3d)**. We also observed that within the *V. natriegens* genome are tRNAs with moderate gene copy number (4-6 genes) but account for less than 1% of the total tRNA pool. These include tRNA^Glu^_UUC_ (0.37%, 6 copies), tRNA^Phe^_GAA_ (0.34%, 4 copies), tRNA^Asn^_GUU_ (0.33%, 5 copies), and tRNA^Cys^_GCA_ (0.43%, 4 copies). tRNA^Gly^ abundance is also lowly abundant despite its high gene copy number (1.65%, 9 copies), especially when compared to *E. coli* strains (∼3%, 4 copies). Intriguingly, previous work has shown the abundances of tRNA^Glu^, tRNA^Cys^, and tRNA^Gly^ are reduced in stressed yeast cells^47^. These tRNAs correspond to the amino acids required for glutathione biosynthesis, and authors propose a mechanism where reductions in these tRNAs can free up the amino acid precursors for glutathione production under conditions of stress^47^. Our culturing conditions for tRNA extraction matched those used for S30 lysate preparation **(see Experimental Section)**, where cells are grown to early to mid exponential phase^31^. However, in *V. natriegens,* these tRNAs may be depleted to compensate for the oxidative stress of rapid aerobic growth^66^.

### tRNA-inclusive codon optimization (TICO) improves translation yield and rate in PURE

With an improved understanding of strain tRNA preferences, we considered whether we could improve translation yields in PURE by better tuning codons to match tRNA abundances. We were particularly interested in addressing the reduced yields observed with PURE translation reactions utilizing Rosetta and BL21 tRNA pools. *In vivo*, codon optimization efforts aim to divert translation resources to the synthesis of a desired protein, often at the expense of the synthesis of other protein products^67^. This is usually done by matching synonymous codons to the preferred codon usage of the organism, resulting in an mRNA sequence highly enriched for codons which are decoded by the most abundant tRNAs^61^. It is unclear whether these or similar codon optimization strategies could improve *in vitro* protein synthesis. Codon optimization attempts in lysate systems have had limited success^29,68^. In a PURE CFE that used *E. coli* W strain tRNAs, optimization to *E. coli* strain W codon usage bias resulted in reduced reaction yields^32^.

Previous work has suggested that translation elongation is rate limiting in CFE^68,69^, and the availability and dynamics of ternary complex sampling on elongating ribosomes appear to be particularly important^69,70^. Most applications of CFE translate a single mRNA into a single gene product, resulting in a narrow subset of tRNAs that are needed for translation. Hence, tRNAs that would otherwise be engaged in the translation of the broader proteome *in vivo* are left underutilized in single-gene CFE. If ternary complex sampling limits translation rates, unproductive ternary complexes harboring underutilized tRNAs could accumulate, competing for ribosome sampling, and would ultimately slow translation. We wondered whether the coding sequence of pJL1 *E. coli* sfGFP may be overly enriched for abundant tRNAs, and would lead to an underutilized tRNA pool that would inhibit its translation. This would be particularly detrimental to tRNA pools like BL21 and Rosetta which have outsized abundances of tRNA^Pro^ _GGG_ and tRNA^eMet^.

To test this hypothesis, we developed a simple codon optimization strategy which aligns codon usage to tRNA abundance. Here, the amino acids within a given protein sequence are assigned to their tRNA isoacceptors according to the relative abundances of individual tRNAs within the isoacceptor family **(Figure 4a, see also Experimental Section)**. The resulting synonymous codon sets for each amino acid are randomly distributed throughout the coding sequence to avoid bias. To further simplify the design, codons were designed to be Watson and Crick cognate matches to the anticodons of their respective tRNAs. To prevent the inclusion of exceptionally rare tRNAs, we excluded tRNAs which encompass less than 10% of the relative abundance within their isoacceptor groups. This tRNA-inclusive codon optimization or TICO strategy aims to engage the majority of the tRNA pool, preventing the accumulation of unproductive ternary complexes that might impede translation. We applied this codon optimization strategy to the abundances determined from tRNA-seq for A19, BL21, and Rosetta tRNA pools generating three sfGFP constructs with subtle changes in synonymous codon usage to each other, but substantial deviations from pJL1 *E. coli* sfGFP **(Supplementary Information SF5)**.

**Figure 4.**
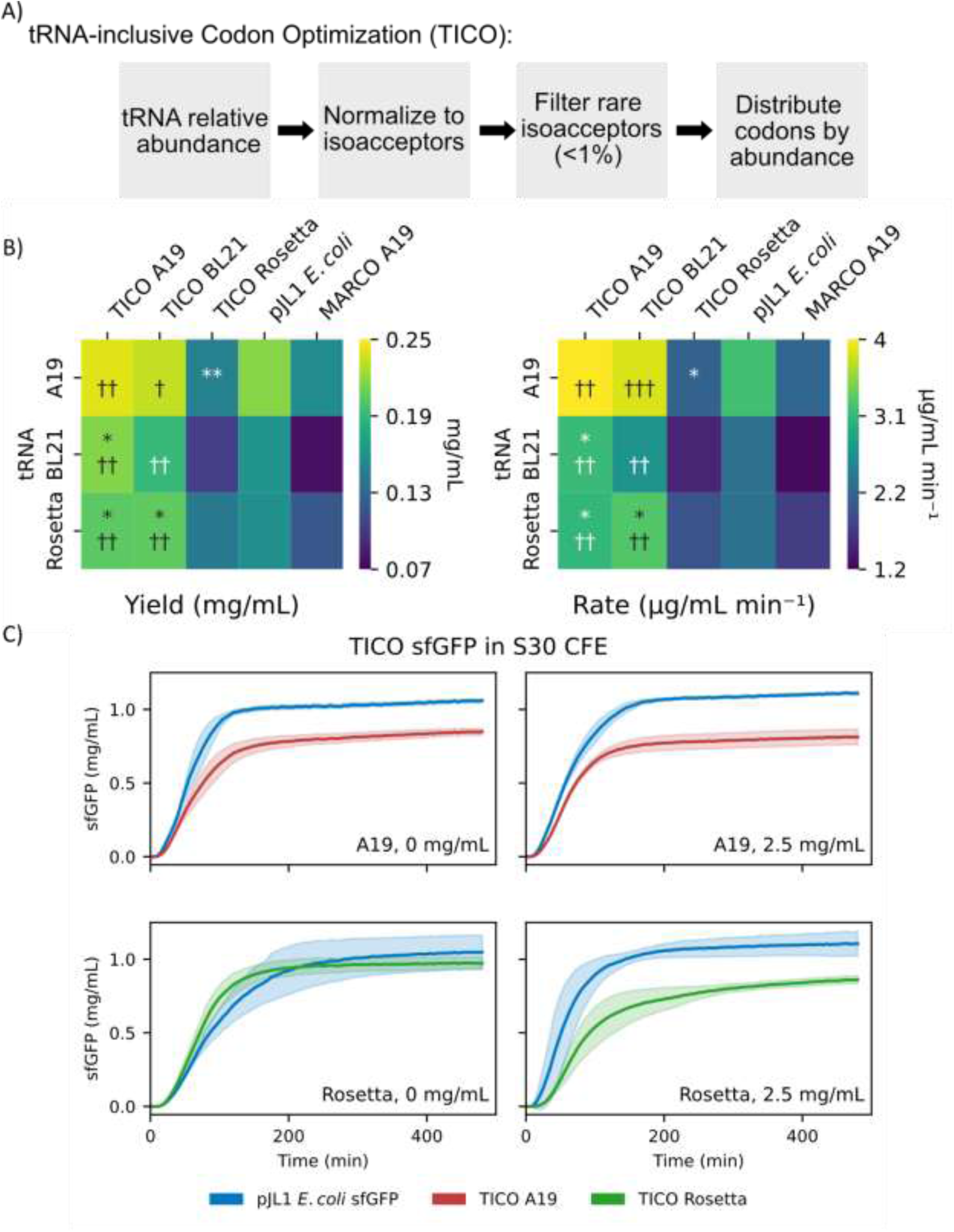
TICO designed sfGFP constructs. **tRNA-inclusive Codon Optimization strategy improves PURE CFE yields relative to pJL1 *E. coli* sfGFP.A)** The TICO design strategy utilizes tRNA relative abundances for each tRNA species and normalizes tRNA species within tRNA isoacceptor families for each amino acid. Normalization is performed by dividing percent abundances of each isoacceptor of an amino acid against the summed percent abundances of the isoacceptor family. Rare tRNAs occupying less than 1% of the isoacceptor family are filtered out while the remaining tRNAs are distributed throughout the coding sequence of the gene of interest approximately to their normalized and filtered abundances. In contrast to TICO, MARCO assigns codons based on the most abundant tRNA among isoacceptors for a given amino acid and serves as a control where tRNA exclusivity is maximized. **B)** Heatmaps of translation yields (left) and translation rates (right) of ΔtRNA PURE completed with either A19, Rosetta, or BL21 tRNA pools translating TICO designed sfGFP constructs. As a control, each tRNA source was tasked with the translation of pJL1 *E. coli* sfGFP. Asterisks represent statistically significant differences in translation for each tRNA against pJL1 *E. coli* sfGFP (* = p < 0.05, ** = p < 0.01) As an additional control, each tRNA source was tasked with the translation of MARCO A19 sfGFP. Daggers represent statistically significant differences in translation for each tRNA against MARCO A19 sfGFP († = p < 0.05, †† = p < 0.01, ††† = p < 0.001). Statistical significance was calculated using two-sample Welch’s t-test. Translation rates were determined by calculating the slope of linear regression of data points within 20-60 minutes of the reaction. **C)** Translation of TICO designed A19 and Rosetta sfGFP constructs within their respective lysates. Top, A19 S30 sfGFP translating either pJL1 *E. coli* sfGFP (blue traces) or TICO A19 (orange traces). Reactions were performed with either no supplemental (left graph) or 2.5 mg/mL of exogenous A19 tRNAs (right graph). Translation of TICO designed A19 and Rosetta sfGFP constructs within their respective lysates. Bottom, A19 S30 sfGFP translating either pJL1 *E. coli* sfGFP (blue traces) or TICO Rosetta(green traces). Reactions were performed with either no supplemental (left graph) or 2.5 mg/mL of exogenous Rosetta tRNAs (right graph). Shaded regions represent standard deviation of mean of three experimental replicates.

*E. coli* pJL1 sfGFP represents a control for standard codon optimization methods, but we envisioned another control that would further test the hypothesis that underutilized tRNAs are inhibitory to translation. If a TICO design strategy aims to be inclusive of tRNAs according to their abundance, a tRNA-*exclusive* strategy which draws only from the most abundant tRNAs should result in the greatest population of underutilized ternary complexes while still engaging an ostensibly optimal tRNA pool. To this end, we designed an sfGFP construct to exclusively engage only the most abundant tRNAs within A19 tRNA, encouraging underutilized tRNAs to accumulate. Because of this, we expected this construct would result in reduced translation yield across all PURE reactions tRNA compared to TICO sfGFPs with Rosetta and BL21 being considerably challenged. We called this design choice Most Abundant tRNA Codon Optimization or MARCO. The MARCO construct has the added benefit of accounting for the impact of designing coding sequences which contain only Watson and Crick cognate matches. It has been previously reported that Watson and Crick cognate matches are more rapidly decoded than wobble counterparts^71^. Notably, MARCO designs for BL21 and Rosetta would be identical to A19 except for the exchange of proline codon CCG to CCC. Since proline accounts for only a small fraction of amino acids of sfGFP (∼4%), the difference in A19 MARCO and BL21/Rosetta MARCO sfGFP translation is expected to be negligible.

TICO had varying effects in improving both translation yield and rate within PURE CFE **(Figure 4b, Supplementary Figure SF6)**. Across all tRNA pools, TICO-designed A19 sfGFP improved both translation rates and end-point translation yields relative to translation of pJL1 sfGFP with BL21 and Rosetta pools benefiting from statistically significant increases in translation yields (p-values of 0.041 and 0.026, respectively, **Figure 4b**). These results suggest that for imbalanced tRNA pools such as BL21 and Rosetta, higher yields may be achieved by engaging with a broader tRNA pool. A similar result was observed for TICO BL21 sfGFP which generated a significant yield improvement for Rosetta tRNA PURE compared to pJL1 *E. coli* sfGFP (p-value = 0.016), and a modest but insignificant improvement for the BL21 and A19 pools compared to pJL1 *E. coli* sfGFP. In contrast and to our surprise, TICO-designed Rosetta sfGFP suffered losses in translation compared to pJL1 *E. coli* sfGFP with the BL21 tRNA pool PURE especially struggling. Still, optimization to Rosetta equalized the yields for A19 tRNA and Rosetta tRNAs.

When translating MARCO A19 sfGFP, translation yields were reduced relative to both TICO sfGFP and pJL1 *E. coli* translation for all tRNA pools (**Figure 4b**). As expected, PURE reactions utilizing either BL21 or Rosetta tRNA pools had a more marked reduction in translation yield compared to the A19 tRNA pool. Surprisingly, BL21 tRNA complemented PURE reactions suffered a greater reduction in translation yield compared to Rosetta tRNA PURE CFE, despite Rosetta tRNAs harboring greater relative abundances of both tRNA^Pro^_GGG_ and tRNA^eMet^ compared to BL21. Still, these results indicate that altering codon preferences to engage in a reduced subset of the tRNA pool negatively impacts translation yields.

We considered if TICO sfGFP constructs would similarly improve translation in lysate-based CFE. To test this hypothesis, we assembled S30 CFE reactions from either A19 or Rosetta lysates and tasked each to translate their respective TICO sfGFPs. In contrast to the results in PURE, TICO sfGFP constructs failed to improve translation yields. Instead, A19 S30 CFE suffered a 19.8% ± 2.3% reduction in yield when translating TICO A19 sfGFP compared to pJL1 *E. coli* sfGFP **(Figure 4b)**. Interestingly, Rosetta S30 was insulated from a similar loss in translation yield with TICO Rosetta sfGFP effectively matching pJL1 *E. coli* sfGFP yields. We hypothesized that reductions in yield may stem from the increased demand of tRNAs that, when combined with the high productivity of S30 CFE, might result in issues in tRNA scarcity. This would mimic our previous observation where insufficient tRNA supply compromised *S. cerevisiae* translation in A19 lysates **(Figure 2c).** To address this possibility, we supplemented exogenous tRNAs to their respective S30 CFE systems to the previously observed saturating final concentration of 2.5 mg/mL. Exogenous tRNAs failed to rescue translation yields of both TICO-designed constructs. In fact, exogenous Rosetta tRNAs reduced TICO Rosetta sfGFP translation yields similar to TICO A19 sfGFP **(Figure 4c)**. From these experiments, we conclude that lysate-CFE is not limited to ternary complex usage.

### *V. natriegens* S30 is not limited by either tRNA abundance or codon optimality

We also wondered if TICO might aid in CFEs which utilize tRNAs harvested from *V. natriegens*. Since the tRNA abundances of *V. natriegens* are quite dissimilar to *E. coli* (**Figure 3a**), we expected that a TICO-informed sfGFP might improve translation within both PURE CFE and *V. natriegens* lysate-based CFE. To test this hypothesis, we assembled PURE CFE reactions where *V*. *natriegens* tRNA pools were used to translate either a TICO-generated *V. natriegens* sfGFP or pJL1 *E. coli* sfGFP. As an additional control, we employed a commercial codon optimization service provided by Integrated DNA Technologies (IDT), generating an sfGFP construct optimized using traditional optimization approaches. While *V. natriegens* and *E. coli* tRNAs differ in tRNA abundances **(Figure 3a)**, pairwise sequence alignments reveal high sequence similarities between their tRNAs, suggesting that they could interchangeably serve as the tRNA complement in *E. coli-*based PURE CFE **(Supplementary Figure SF7).** Still, we prepared an additional control where the *V. natriegens* sfGFP constructs are translated by PURE CFE reactions using A19 tRNA pools.

*V. natriegens* tRNAs struggled to translate pJL1 *E. coli* sfGFP, TICO-generated *V. natriegens*, and commercially optimized *V. natrigens* sfGFP compared to reactions containing A19 tRNAs (**Figure 5a**). Surprisingly, IDT-optimized *V. natriegens* sfGFP translated well with the A19 tRNAs, increasing translation rates compared to pJL1 *E. coli* sfGFP and achieved similar end-point yields. PURE CFE using A19 tRNAs translated *V. natriegens* sfGFP with a ∼40% greater yield compared to reactions which used *V. natriegens* tRNAs **(Figure 5a)**. TICO *V. natriegens* sfGFP was similarly poorly translated by *V. natriegens* tRNAs. These results suggest a limited compatibility of *V. natriegens* tRNAs with the *E. coli* translational machinery within the PURE CFE, despite the high sequence similarity. For PURE reactions completed with *V. natriegens* tRNAs, both IDT and TICO designed *V. natriegens* sfGFP constructs outperformed pJL1 *E. coli* sfGFP in translation, indicating that matching to tRNA is still critical despite the inherently reduced translation. Unfortunately, TICO *V. natriegens* sfGFP failed to improve yields relative to commercially optimized *V. natriegens* tRNAs, contrary to our observations with the *E. coli* TICO constructs.

**Figure 5.**
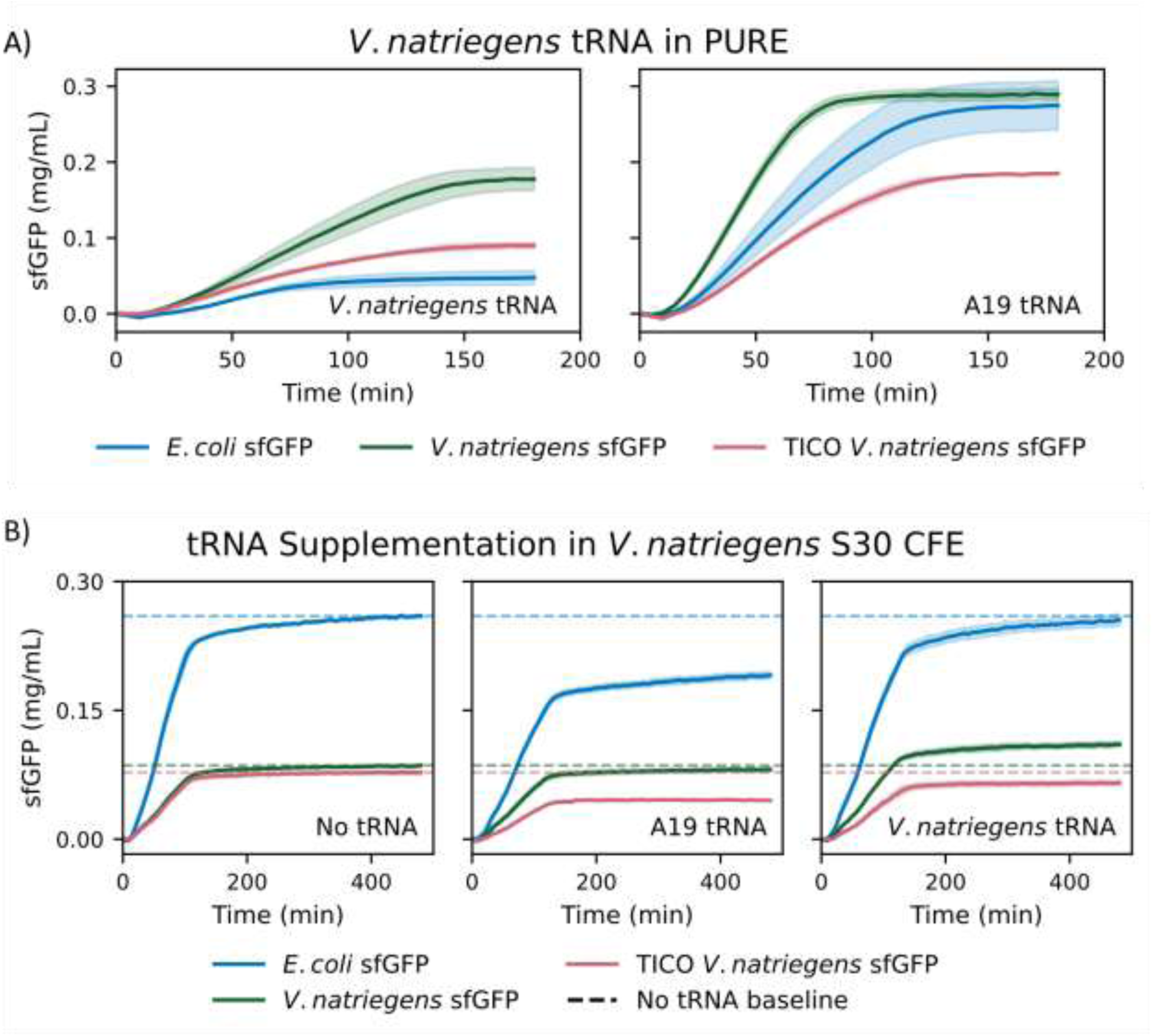
*V. natriegens* and *E. coli* translation machinery have limited mutual compatibility. *V. natriegens* and *E. coli* translation components have limited compatibility within PURE or S30 CFEs. **A)** *V. natriegens* tRNA (left) or A19 tRNA (right) completed PURE CFE reactions translating either pJL1 E. coli sfGFP (blue traces), commercially optimized *V. natriegens* sfGFP (green traces), or TICO *V. natriegens* sfGFP (pink traces). **B)** *V. natriegens* S30 translating pJL1 E. coli sfGFP, commercially optimized *V. natriegens* sfGFP, TICO V. natriegens sfGFP with either A19 or *V. natriegens* tRNA supplemental tRNAs. Reactions contained either no supplemental tRNA (left), 2.5 mg/mL added A19 tRNA (center), or 2.5 mg/mL *V. natriegens* tRNA (right). Colored dashed lines note end-point fluorescence for *V. natriegens* S30 CFE reactions containing no supplemental tRNAs across sfGFP plasmids. Shaded regions represent standard deviation of mean for three experimental replicates.

*V. natriegens* lysates have been used in S30 CFE systems, but their yields lag behind traditional *E. coli* lysate-based CFEs^29,54,72^. When developing new S30 systems, it is common practice to supplement non-*E. coli* lysates with *E. coli* MRE600 tRNAs during reaction optimization^48^. We were interested if adding supplemental, native *V. natriegens* tRNAs might improve yield of its own CFE, especially given the limited compatibility observed between *V. natriegens* tRNAs with *E. coli* translation machinery. To test this, we assembled *V. natriegens* S30 CFEs tasked with the translation of either pJL1 *E. coli* sfGPF, TICO-generated *V. natriegens* sfGFP, or IDT *V. natriegens* sfGFP. Additionally, *V. natriegens* S30 reactions were supplemented with either isolated A19 tRNAs or *V. natriegens* tRNAs again at concentrations considered saturating (2.5 mg/mL) from our previous experiments.

We were surprised to find that the pJL1 *E. coli* sfGFP produced the greatest sfGFP yields despite both TICO-generated *V. natriegens* sfGFP and IDT *V. natriegens* sfGFP performing better in PURE reactions containing *V. natriegens* tRNA **(Figure 5b)**. Although pJL1 *E. coli* sfGFP was the best translating plasmid within the lysate CFE, supplemental A19 tRNA failed to improve translation yields. In fact, both TICO *V. natriegens* sfGFP and pJL1 *E. coli* sfGFP had markedly reduced yields when supplemented with A19 tRNAs. In contrast, supplementing *V. natriegens* S30 reactions with *V. natriegens* tRNA had little effect on translation for pJL1 *E. coli* sfGFP and only modestly improved the translation of IDT *V. natriegens* sfGFP **(Figure 5b)**. Given the inhibitory effect of A19 tRNA within *V. natriegens* S30 CFE and the reduced yields from *V. natriegens* tRNA in *E. coli* PURE, we conclude that there is a mutual incompatibility between *E. coli* and *V. natriegens* translation machinery. It may be that the tRNAs of each organism are similar enough to engage with each others’ translational elements but cannot be simply substituted, effectively resulting in mutual competitive inhibition. Moreover, our results suggest that *V. natriegens* S30 is not limited by tRNA abundance nor codon choice, running counter to our initial hypothesis and our observations of PURE CFE.

## Discussion

In this study, we found that small RNA isolation as described by Avcilar-Kucugoze et al.^43^ generates tRNAs capable of translation within PURE and S30 CFEs. We established *E. coli* A19 as a reliable source of tRNAs for PURE CFE, resolving a critical reagent limitation within the cell-free translation field. We also found tRNAs sourced from either BL21 or Rosetta exhibited reduced activity in PURE. We later found that BL21 and Rosetta tRNA contain an outsized abundances of certain tRNAs such as tRNA^Pro^_GGG_ and tRNA^eMet^_CAU_. We were also able to mitigate the reduced translation of sfGFP from these pools through codon design, called TICO, by engaging a broader subset of the tRNA pool.

Efforts in enhancing the translational capabilities of PURE have largely centered on improvements within the protein components of PURE either by supplementation of additional components or increasing their general abundances^73^. These additive strategies can conflict with the goal of maintaining a minimal translation system, as each new element increases complexity and introduces potential sources of variability. Codon optimization represents an attractive solution to improving PURE CFE translation yields while maintaining its minimalist approach.

Previous work identified ternary complex sampling on the elongating ribosome as a critical determinant of translation efficiency, and tuning tRNA pools to codon demands reduced the latency of elongation events, increasing translation speed^70^. TICO supports these observations by leveraging a broader range of tRNAs, enhancing both translation rate and yield. Similarly, when a greater abundance of tRNAs were excluded through codon design, translation rates and yields suffered. Previous work explored the complete minimization of elongation latencies using the *de novo* synthesis and reassembly of tRNA pools tuned to coding sequences^70^. TICO similarly achieves this by repurposing a tRNA pool evolved for *in vivo* translation to better serve single-gene *in vitro* expression.

The TICO strategy described in this study represents a simplified approach to PURE codon optimization, and our preliminary examinations here were designed to isolate codon effects without accounting for other determinants of translation such as mRNA secondary structure^74,75^, wobble decoding^71^, or internal Shine-Delgarno sequences^76^. Further refinements to TICO which consider these factors are expected to improve translation efficiency further still. Though useful for single gene CFE, we expect TICO to become less impactful as a greater number of genes are introduced to PURE CFE. More nuanced approaches will have to contend with mutually shared tRNA demands across a broader proteome^67^. This will become especially important in the development of a self-replicating PURE system which must balance assembly of a minimal tRNA pool that can still meet the challenge of homeostasis and replication. Still, in PURE CFE, our current TICO approach represents a rare improvement in translation yield through codon optimization where other efforts have failed^32^.

We were unable to improve the translation yields of either *E. coli* or *V. natriegens* S30 CFE. Exogenous tRNAs were unnecessary in *E. coli* S30 CFE unless codon choice shifted demand to tRNAs underrepresented in the cytosol. In *V. natriegens*, exogenous tRNAs were also unnecessary and if *E. coli* tRNAs were used, were inhibitory to translation. Similarly, TICO-optimized constructs failed to improve translation for both *E. coli* and *V. natriegens* S30 CFE. These observations suggest that CFE limitations likely arise from factors beyond the core translation machinery which is likely already optimized for maximal protein expression^77^. Indeed, *V. natriegens* CFE boasts the highest yields among non-*E. coli* prokaryotic CFE systems, perhaps a consequence of its robust translation machinery needed to enable its fast doubling time. *V. natriegens* supports abundant ribosome synthesis, generating an estimated 115,000 ribosomes per cell, compared to 70,000 in *E. coli*^78^. Notably, we found that *V. natriegens* has a significantly higher fraction of initiator tRNA in its tRNA pool compared to *E. coli* strains (∼15% vs 3-4%). Initiator tRNAs are involved with ribosome synthesis and assembly^79,80^. Thus, *V. natriegens* may appropriate additional initiator tRNAs to support its increased ribosome biogenesis. In summary, the inherent synthesis capabilities of *V. natriegens* S30 CFE appear to render supplemental tRNAs unnecessary. Still, exogenously supplied, native tRNAs might improve less robust lysate-based CFE systems as was previously observed with Syn3A CFE^81^. We expect weakly translating lysates with poorer shared sequence similarity to *E. coli* tRNAs **(Supplementary Figure SF7)** to be best positioned to benefit from native exogenous tRNA pools.

In this study, we considered a narrow scope of *E. coli* strains as tRNA sources for CFE because of their popularity in lysate-based CFE systems. Further improvements in PURE CFE yields could be realized from tRNA pools sourced from strains more competent for generalized protein expression. For example, we show that the pRARE2 plasmid does reshape the Rosetta tRNA pool but in a way which exacerbates a mismatch between tRNA supply and demand for sfGFP translation. Intelligent design of tRNA expression plasmids which better address codon bias might further improve translation yields in PURE CFEs^70^. We expect a combination of tRNA-seq and TICO will be critical in these developments, and the work described here presents a roadmap for these efforts.

### Conclusions

We have applied a method to isolate tRNAs capable of serving as substrates for both PURE and S30 CFE. We find that *E. coli* A19 tRNAs match the translation yields of both kit-provided and the now defunct commercial Roche MRE600 tRNAs. We also find exogenous tRNAs are generally unnecessary for S30 CFE systems. Our efforts address a critical reagent challenge for users of CFE and highlight the importance of tRNA abundance and codon bias in PURE protein expression.

## Experimental Section

### Isolation of *E. coli* tRNAs

Methods for *E. coli* tRNA isolation were used as described by Avcilar-Kucugoze et al.^43^ with some modifications. For each strain of *E. coli*, overnight cultures were used to inoculate 1L of Terrific Broth. In the case of Rosetta2 (DE3), cells were grown in the presence of 34 µg/ml chloramphenicol to maintain the pRARE2 plasmid. Cultures were grown to an OD_600_ of ∼2.0 (∼3 hrs) and harvested by centrifugation at 4,500 xg for 20 minutes at 4°C. Cells were washed once with 0.9% NaCl, spun again at 4,500 xg for 20 minutes at 4°C. The wash solution was discarded and the resulting pellets were weighed and stored at - 80°C until further processing.

Frozen cell pellets were resuspended in 18 mLs of Extraction Buffer (50 mM sodium acetate, 10 mM magnesium acetate, pH 5.0) by vortexing. To the resuspended solution, an equal volume of acid-buffered phenol at pH 4.5 (VWR International, Radnor, PA, USA, catalog number: 97064-716 or Invitrogen, Thermo Fisher Scientific, Waltham, MA, USA, catalog number: AM9722) was added and the resulting solution was mixed in a shaking incubator for 30 minutes at 37°C. The emulsion was then separated by centrifugation at 4,500xg for 15 minutes at 4°C. The aqueous layer was collected and an additional 14 mL of Extraction Buffer was added. The solution was mixed as before for 15 min before being centrifuged again at 4,500xg for 15 minutes at 4°C. Again, the top layer collected and added to the previously collected aqueous phase.

To the collected aqueous phases, 5M NaCl was added to achieve a final concentration of 0.2 M NaCl to precipitate rRNAs. The solution was briefly mixed by repeated inversion and an equal volume of isopropanol was added to begin precipitation. The solution was briefly mixed by inversion and was then centrifuged at 14,500xg for 15 min at room temperature. The resulting pellet was rinsed with ice-cold 70% ethanol and allowed to air-dry. The pellet was then resuspended in 15 mL of cold 1M NaCl and centrifuged at 9,500xg for 20 min at 4°C.

The supernatant was collected, and 30 mLs of cold ethanol was added to precipitate the remaining nucleic acids (tRNA and DNA). The solution was allowed to precipitate for at least 30 minutes at −20C before being centrifuged at 14,500xg for 5 min at 4°C. The pellet was rinsed with ice-cold 70% ethanol and allowed to air-dry.

The resulting tRNA and DNA pellet was resuspended in 6 mL of 300 mM sodium acetate pH 5.0 and 2.3 mL of isopropanol was added to the solution. After incubating for 10 minutes, the solution was centrifuged at 14,500xg for 5 minutes at room temperature. The supernatant was recovered, and an additional 3.4 mL of isopropanol was added to precipitate the tRNAs. After incubating for at least 30 minutes at −20°C, the solution was centrifuged at 14,500xg for 15 minutes at 4°C. The pellet was air-dried and resuspended in 400 µL of ddH2O. The concentration was determined from A_260_ using a Nanodrop ND-1000 spectrophotometer (Thermo Fisher Scientific), aliquoted, and stored in the −80°C until needed.

### Isolation of *V. natriegens* tRNA

To isolate *V. natriegens* tRNAs, the same isolation procedure for *E. coli* tRNAs was performed but with the following modifications. First, overnight *V. natrigens* cultures were grown at 30°C and used to inoculate 1L of LB-media supplemented with V2 salts (200 mM NaCl, 23.1 mM MgCl_2_, and 4.2 mM KCl). Cultures were grown to an OD_600_ of 1-2 at 37°C before harvesting through centrifugation at 4,500xg for 20 minutes at 4°C. The cells were washed as described above before storage at −80°C until processing.

*V. natriegens* tRNA isolation proceeded as described above except for the separation of DNA and tRNA. For *V. natriegens,* the tRNA and DNA pellet was resuspended by 6 mL of 300 mM sodium acetate pH 5.0 and 0.5 mL of isopropanol was added to precipitate DNA. The resulting solution was pelleted via centrifugation at 14,500xg for 5 minutes at room temperature. The resulting supernatant was recovered and 5.2 mL of isopropanol was added to precipitate the remaining nucleic acids after incubation at −20°C for 30 minutes. The solution was pelleted by centrifugation at 14,500xg for 15 minutes at 4°C. The resulting pellet was resuspended in 10 mL of 300 mL sodium acetate pH 5.0 and 6 mL of isopropanol were added to precipitate any remaining DNA. The solution was incubated for 10 minutes at room temperature and pelleted through centrifugation at 14,500xg for 5 minutes at room temperature. The supernatant was again recovered and an additional 4 mL of isopropanol was to precipitate the tRNA. The solution was incubated at −20°C overnight before a final centrifugation step at 14,500 for 15 minutes at 4°C. The resulting pellet was washed once with ice-cold 70% ethanol and allowed to air-dry. The pellet was resuspended in 200 µL and the resulting tRNA bulk pool quantified using a Nanodrop spectrophotometer.

### PURE Cell-Free Expression

PURE CFE reactions were prepared using NEB PURExpress ΔtRNA,ΔAA kits (New England Biolabs, Ipswich, MA, USA). 12.5 µL reactions were assembled with 2.5 µL of Solution A, 1.25 µL amino acid mixture, 3.75 µL of Solution B, 5 nM DNA template, and 25 units of Murine RNAse Inhibitor (New England Biolabs, Ipswich, MA, USA). In-house isolated tRNA pools from either A19, BL21, and Rosetta *E. coli* strains complemented the reactions at final concentration of 3.5 mg/mL. Kit-supplied control tRNA provided by the NEB PURExpress ΔtRNA,ΔAA kit served as the positive control. Where relevant, *E. coli* MRE600 tRNAs from Roche Diagnostics (Roche Diagnostics, Mannheim, Germany, discontinued) were also used as a control. Researchers seeking to replicate this work are invited to contact the corresponding author for MRE600 controls if still available. To account for potential batch-to-batch variability, data for each figure were gathered from their own set of NEB PURExpress ΔtRNA,ΔAA kits. Multiple kits were pooled when necessary.

Assembled PURE CFE reactions translating sfGFP were monitored using SpectraMAX Gemini EM MicroPlate Reader tracking fluorescence emission at 509 nm from excitation at 488 nm, taking readings every 5 minutes for 3 hrs at 37°C. For PURE CFE reactions translating NanoLuc, reactions were incubated at 37°C for 3 hrs. End-point yields were determined using the NanoGlo HiBiT Extracellular Detection System (Promega, Madison, Wisconsin, USA). Briefly, translated reactions were diluted 1:100 into miliQ H_2_O. 10 µL of the resulting dilutions were mixed with an equal volume of 1x NanoGlo HiBiT Extracellular reagent. Luminescence was tracked for 10 minutes using SpectraMAX Gemini EM MicroPlate Reader.

### Generation of A19, Rosetta, and *Vibrio natriegens* Lysates

Lysates for A19 and Rosetta S30 CFE were generated according to Garamella et al^82^. Fresh overnight cultures were used to inoculate 1L of 2XYTPG media. Cultures were grown at 37°C until achieving an OD of 1.0 before being harvested via centrifugation at 4,500xg for 20 minutes at 4°C. In the case of Rosetta, overnight and outgrowth cultures were grown with the inclusion of chloramphenicol. Pelleted cells were washed twice with 30 mL of chilled S30 Extract Buffer (14 mM magnesium acetate, 60 mM potassium acetate, 10 mM Tris acetate, pH 8.2) through resuspension followed by centrifugation at 4,500xg for 20 minutes at 4°C. Washed pellets were flash frozen in liquid nitrogen and stored at −80°C until processed.

Stored cells were then thawed on ice and resuspended in 1 mL of S30 Extract Buffer per gram wet cell mass. Resuspended cells were lysed using sonication using a Sonic Dismembrator (model FB120, probe model no. CL-18, Fisher Scientific) 20kHz at 50% amplitude with 15 seconds on and 10 seconds off achieving a total of ∼2.7 kJ delivered. Cellular debris was cleared by centrifugation at 16,000xg for 30 minutes at 4°C. The supernatant was collected, aliquoted into 500 uL volumes into 15 mL conical tubes before incubation at 37°C for 60 minutes in a run-off step. Following translation run-off, lysates were collected and centrifuged once more at 16,000xg for 30 minutes at 4°C to clear remaining debris. Supernatants were collected, aliquoted, and flash frozen in liquid nitrogen before storage at −80°C.

*V. natriegens* lysates were prepared according to Wiegend et al^31^. Briefly, overnight cultures of *V. natriegens* were used to inoculate 1.5 L of LB media supplemented with V2 salts (200 mM NaCl, 23.1 mM MgCl_2_, and 4.2 mM KCl). Inoculum was grown at 30°C until achieving an OD600 of ∼1.0. Cells were then harvested by centrifugation at 4,500xg for 20 minutes at 4°C. Pelleted cells were washed twice with 30 mL of chilled S30 Extract Buffer (14 mM magnesium acetate, 60 mM potassium acetate, 10 mM Tris acetate, pH 8.2) before flash freezing in liquid nitrogen and storage at −80°C until processing. The pellet was resuspended in 0.8x mL of S30 Extract Buffer, the cells were lysed by sonicating with a frequency of 20kHz at 50% amplitude with 15 seconds on and 10 seconds off achieving a total of ∼1.5 kJ delivered. Cell debris was clarified using centrifugation at 16,000 rpm for 30 minutes at 4°C. Clarified lysates were separated into 50 uL aliquots avoiding a thin top layer, as well as the pellet cell debris.

### S30 Cell-free expression of sfGFP

A19, Rosetta, and *V. natriegens* lysate-based CFE reactions were assembled according to previous studies with some modifications^26,82^. 16 µL reactions were assembled containing final concentrations of 5 nM DNA template, 1.5 µL T7 RNA polymerase, 0.4 U/μL Murine RNase Inhibitor (New England BioLabs), 12 mM magnesium glutamate, 140 mM potassium glutamate, 2 mM amino acid mix, 1.5 mM ATP and GTP, 0.9 mM CTP and UTP, 0.26 mM coenzyme A, 0.33 mM NAD, 0.75 mM cAMP, 0.068 mM folinic acid, 1 mM spermidine, 30 mM 3-PGA, 1 mM DTT, and final reaction volume of 35% of lysate (5.6 μL).

Assembled S30 CFE reactions translating sfGFP were monitored in a clear-bottomed, black 384-well plate using SpectraMAX Gemini EM MicroPlate Reader tracking fluorescence emission at 509 nm from excitation at 488 nm and taking readings every 5 minutes for 8 hrs at 30°C.

### sfGFP Standard Curve Generation

To generate a standard curve for absolute protein yield quantification, a His_6_-tag was inserted into the C-terminus of pJL1 *E. coli* sfGFP using Q5 mutagenesis kit according to manufacturer’s protocol (New England Biolabs, E0554S). Successful transformations were sequenced using whole plasmid sequencing (Plasmidsaurus, San Francisco, CA, USA). His-tagged sfGFP was transformed into *E. coli* BL21 DE(3) for expression. sfGFP was expressed relying on the leaky expression of the pJL1 *E. coli* sfGFP plasmid.

Briefly, 10 mL of overnight cultures were used to inoculate 1L of LB media supplemented with 50 µg/mL kanamycin and were allowed to grow uninduced for 9 hrs. Cells were pelleted using centrifugation at 3,400xg for 20 minutes at 4°C and washed once with 30 mL of 0.9% NaCl before centrifugation again at 3,900xg for 20 minutes at 4°C. The pellets were then flash frozen in liquid nitrogen and stored at −80°C until processing.

The pellet was resuspended by vortexing in 15 mL wash buffer (50 mM HEPES-KOH pH 7.6, 1 M NH4Cl, 10 mM MgCl2, 15 mM Imidazole, 7 mM 2-mercaptoethanol, 1X Halt Protease inhibitor Cocktail (Thermo Scientific)). Resuspended cells were sonicated on ice at 50% power, 15 sec on, 15 sec off until 2 kJ was reached. The lysate was mixed and the lysate was allowed to cool on ice for 5 min before the sonication step was repeated once. The lysate was centrifuged at 15,000 x g for 15 min at 4 °C. The supernatant was saved and mixed with 1 mL of equilibrated (with wash buffer) Ni-NTA agarose bead slurry (Goldbio). This mixture was incubated at 4 °C with rocking for 30 minutes. In a 4 °C cold room, the mixture was poured through a fritted column and the flow through was reapplied to the column once. The resin was washed three times with 7 mL of wash buffer. sfGFP was eluted from the column by applying 2 mL elution buffer (50 mM HEPES-KOH pH 7.6, 100 mM KCl, 10 mM MgCl2, 300 mM Imidazole, 7 mM 2-mercaptoethanol) to the resin and collecting the flow through. The purified sfGFP was dialyzed using a 3K MWCO slide-a-lyzer (Thermo Scientific) against 1 L of storage buffer (100 mM Tris-HCl pH 7.6, 200 mM KCl, 20 mM MgCl2, 5 mM BME, 20% glycerol) at 4 °C. After an overnight dialysis, the slide-a-lyzer was removed and put in a fresh liter of storage buffer and dialyzed for another 6 hours at 4 °C. The purified sfGFP was collected, aliquoted into single use tubes, and snap frozen in liquid nitrogen. The purified protein was stored at −80 °C until use.

Protein concentration of sfGFP was determined using Pierce 660 assay and a standard curve was assembled from a known concentration of sfGFP to observed relative fluorescence units. The resulting standard curve was used to determine absolute concentrations of sfGFP synthesized in CFE reactions.

### Small RNA Sequencing

Total RNA (2.5 μg) was resuspended in 100 mM Tris-HCl pH 9, 2.5 ng of each synthetic tRNA control (Supplementary Table ST1) was added, and the mixture was deacylated at 37°C for 1 hour. Samples were dephosphorylated with 2U rSAP (NEB #M0371S) in 1x rCUTSMART Buffer at 37°C for 30 mins followed by deactivation at 65°C for 5 mins. tRNA-seq libraries were prepared as previously described^64^ with modifications to make the workflow compatible with an automated liquid handler. All oligonucleotides used in this study were obtained from Integrated DNA Technologies (IDT) **(Supplementary Table ST1)**. Both adapters are blocked by the 3′ chain terminator dideoxycytidine to prevent concatemer formation, and 5′-phosphorylated to enable pre-adenylation by Mth RNA ligase prior to ligation^83^. To remove large RNA species (> 200 nt), 37.5 μL of room-temperature-equilibrated SPRISelect beads (Beckman Coulter, B23318), 55 μL isopropanol, and RNAse-free water was added to the reaction to make up a final volume to 105 μL, pipetting gently up and down, and incubated for 5 minutes at room temperature. The bead mixture was incubated on a magnetic rack and the supernatant containing the small RNA was transferred to a new tube. An additional 37.5 μL SPRISelect beads, 55 μL isopropanol, and RNAse-free water were added to the supernatant to make up the final volume to 220 μL, pipetting gently up and down, and incubated for 5 minutes at room temperature on a Hula Mixer. The beads were washed with freshly prepared 70% ethanol and left to air dry. The samples were eluted by resuspending the beads in nuclease-free water and incubating them for 10 minutes at room temperature. The RNA concentration was determined using RNA HS Qubit Fluorometric Quantification. Next, ligation was performed for 2 hours at 25°C in a 20 μL reaction volume containing pre-adenylated 3’ adapter and small RNA substrate in a 4:1 molar ratio, 1x T4 RNA Ligase Reaction Buffer, 200 U of T4 RNA ligase 2 (truncated KQ; NEB), 25% PEG 8000, and 10 U SUPERase In (Ambion). To size select the ligation product, 35 μL of room-temperature-equilibrated SPRISelect beads (Beckman Coulter, B23318), 48 μL isopropanol, and RNAse-free water was added to the reaction to make up a final volume to 160 μL, pipetting gently up and down, and incubated for 5 minutes at room temperature. The beads were washed with freshly prepared 70% ethanol and left to air dry. The samples were eluted by resuspending the beads in nuclease-free water and incubating them for 10 minutes at room temperature. For primer-dependent reverse transcription reactions, adapter-ligated tRNA, RT primer, and 0.5 mM dNTPs were added to a PCR reaction tube, denatured at 95°C for 2 mins and annealed at 60°C for 30 sec and then cooled to 4°C at the lowest ramp rate (0.2°C/s) in a Thermocycler. Reverse transcription was performed in the presence of 1X Maxima RT Buffer, 10 U SUPERase In, and 200 U Maxima RT (Thermo Scientific #EP0742) at 55°C for 60 mins. The reaction was at 85 °C for 5 mins and cooled to 4 °C. Template RNA was subsequently hydrolyzed by the addition of 1 μl 5M NaOH and incubation at 95°C for 3 mins. To purify the single-stranded DNA ligation product, 35 μL of room-temperature-equilibrated SPRISelect beads (Beckman Coulter, B23318), 48 μL isopropanol, and RNAse-free water was added to the reaction to make up a final volume to 160 μL, pipetting gently up and down, and incubated for 5 minutes at room temperature. The beads were washed with freshly prepared 70% ethanol and left to air dry. The samples were eluted by resuspending the beads in nuclease-free water and incubating them for 10 minutes at room temperature. PCR was used to attach Illumina P7/P5 sequences to flank the tRNA insert. Each PCR was set up to contain 5 μL ssDNA ligation product, 25 μL 2X NEBNext Ultra II Q5 Master Mix, and 20 μL water. The PCRs were incubated at 98°C for 30 s followed by 12 cycles of 98°C for 10 s and 65°C for 75 s, and a final extension at 65°C for 5 m. Final amplified libraries were purified via SPRISelect beads using the manufacturer’s protocol. Individual libraries were quantified using Qubit dsDNA BR Assay Kit following manufacturer’s protocol, pooled at equimolar ratio. Finally, the pooled barcoded tRNA-seq libraries were sent to the Biopolymers Facility (Harvard Medical School) for Full Lane NovaSeq X Plus 10B sequencing.

A curated tRNA reference library was constructed to enable accurate quantification of tRNAs across Escherichia coli and *Vibrio natriegens* strains. All tRNA sequences for *E. coli* K-12 MG1655 and BL21-DE3 were downloaded from GtRNAdb (accessed 3rd April, 2025). For organisms or plasmids not represented in GtRNAdb, such as *V. natriegens* (NCBI NZ_CP009977.1) and the pRARE2 expression plasmid (Addgene #62651), mature tRNA genes were predicted using tRNAscan-SE v2.0^84^ in bacterial mode. All tRNA sequences were processed with custom Python scripts to remove introns, append terminal CCA tails where missing, and collapse redundant sequences. Each retained sequence was assigned a unique identifier based on its source and anticodon. When identical sequences were present in both *E. coli* and *V. natriegens*, they were retained and annotated to reflect their shared identity, since the respective samples were processed independently. The resulting reference contains 91 non-redundant tRNA sequences with standardized FASTA headers encoding tRNA identity, source organism, anticodon, and duplication status. Finally, both synthetic tRNA reference sequences **(Supplementary Table ST1)** which were used as experimental controls were added to the reference dataset used in all downstream alignment and quantification steps. Demultiplexed tRNA-seq reads were processed and aligned as previously described^85^ using the boilerplate example of the whole read processing workflow on GitHub with minor modifications: https://github.com/krdav/tRNA-charge-seq/blob/main/projects/example/process_data.ipynb. Briefly, read pairs were trimmed and merged using AdapterRemoval using i7/i5 index sequences defined for the given sample and minimum length set to 25. Each merged reads file was split based on the adaptor barcode (we only used 1 barcode for all samples to reduce sequencing bias) and the sequence was trimmed off the 3’ end; similarly, the 0-4 nt. staggered UMI was located, saved, and trimmed off the 5’ end, leaving only the tRNA sequence with possible 5’ non-template bases introduced during RT-PCR. Trimmed reads were aligned to a masked reference as described below using the Smith-Waterman algorithm implemented by SWIPE^86^. We defined the input score matrix: a match score of 1, a mismatch score of –3, and a score for alignment to a masked reference position (N) of 0. To analyse the tRNA-seq data, we extracted count data from the alignment results and calculated RPM (reads per million) for each tRNA annotation.

### TICO and MARCO Design Strategy

TICO-designed sfGFP coding sequences were designed using an in-house Python program. For each tRNA pool, individual tRNAs were grouped by their amino acid isoacceptor class and assigned weights based on their percent relative abundance within the pool. Within each isoacceptor group, tRNA abundances were normalized to the total tRNA abundances for that amino acid. To avoid biasing codon usage toward exceedingly rare tRNAs, isoacceptors representing less than 10% of the total abundance (weights less than 0.1) for their class were excluded. Codon assignments for each amino acid were then made by randomly selecting from the remaining isoacceptors, proportionally to their normalized weights, and applied sequentially across the sfGFP coding sequence. A similar program was used to generate MARCO A19, but rather than assign codons proportionally to normalized weights, codons were assigned based on the most abundant tRNA isoacceptor within the amino acid class.

## Data Availability

Raw tRNA sequencing data have been deposited in the Sequence Read Archive (SRA) under accession PRJNA1285607.

## Acknowledgments

This work was supported by Alfred P. Sloan Foundation grant G-2024-22710 and NSF awards 2419641 (to K.P.A.), NSF award 2338121 (to A.E.E.), and US Department of Energy (DOE) grant DE-FG02-02ER63445, and NSF Award 2123243 (both to G.M.C.).

## Supplementary Information

**Supplementary Table 1.**
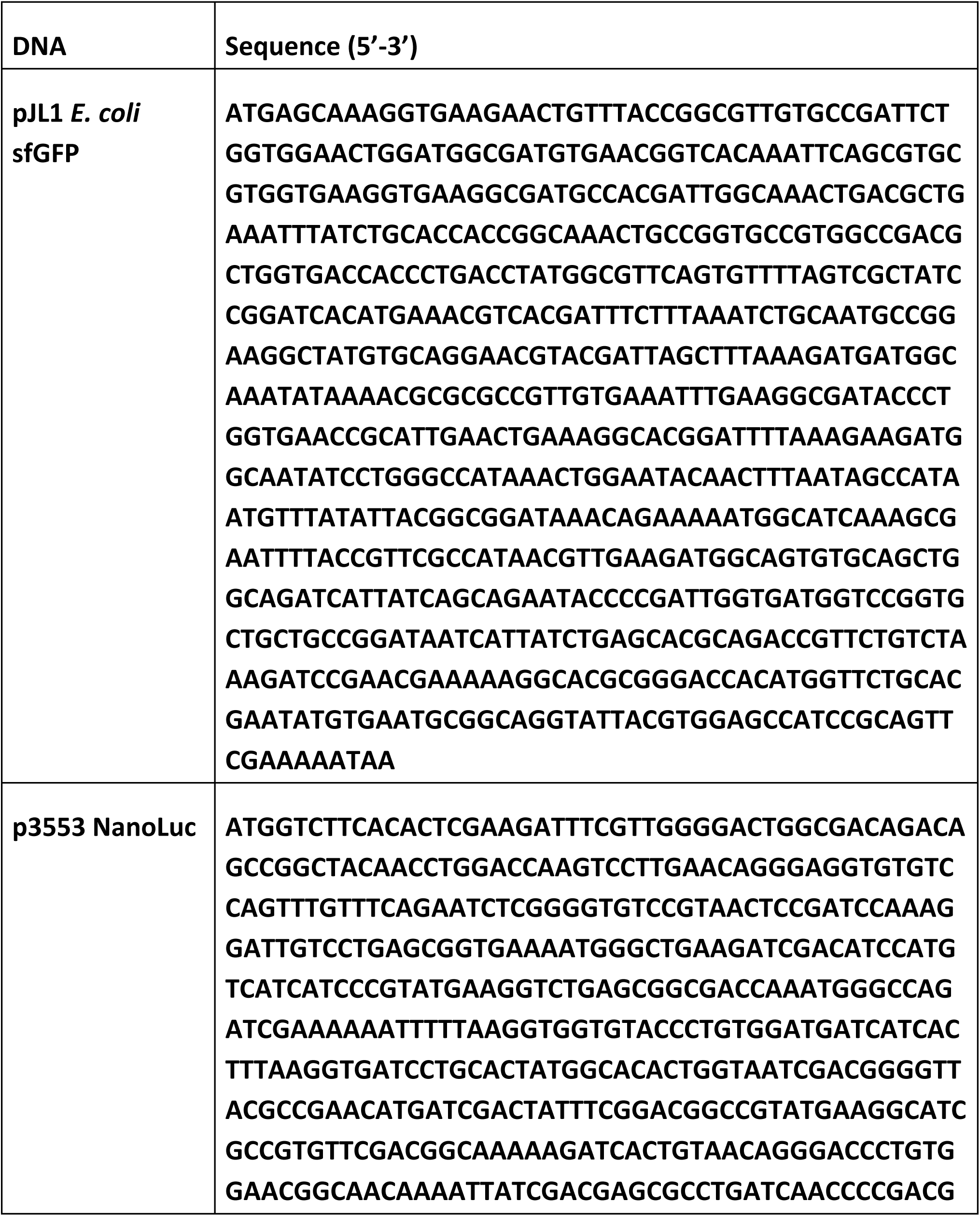
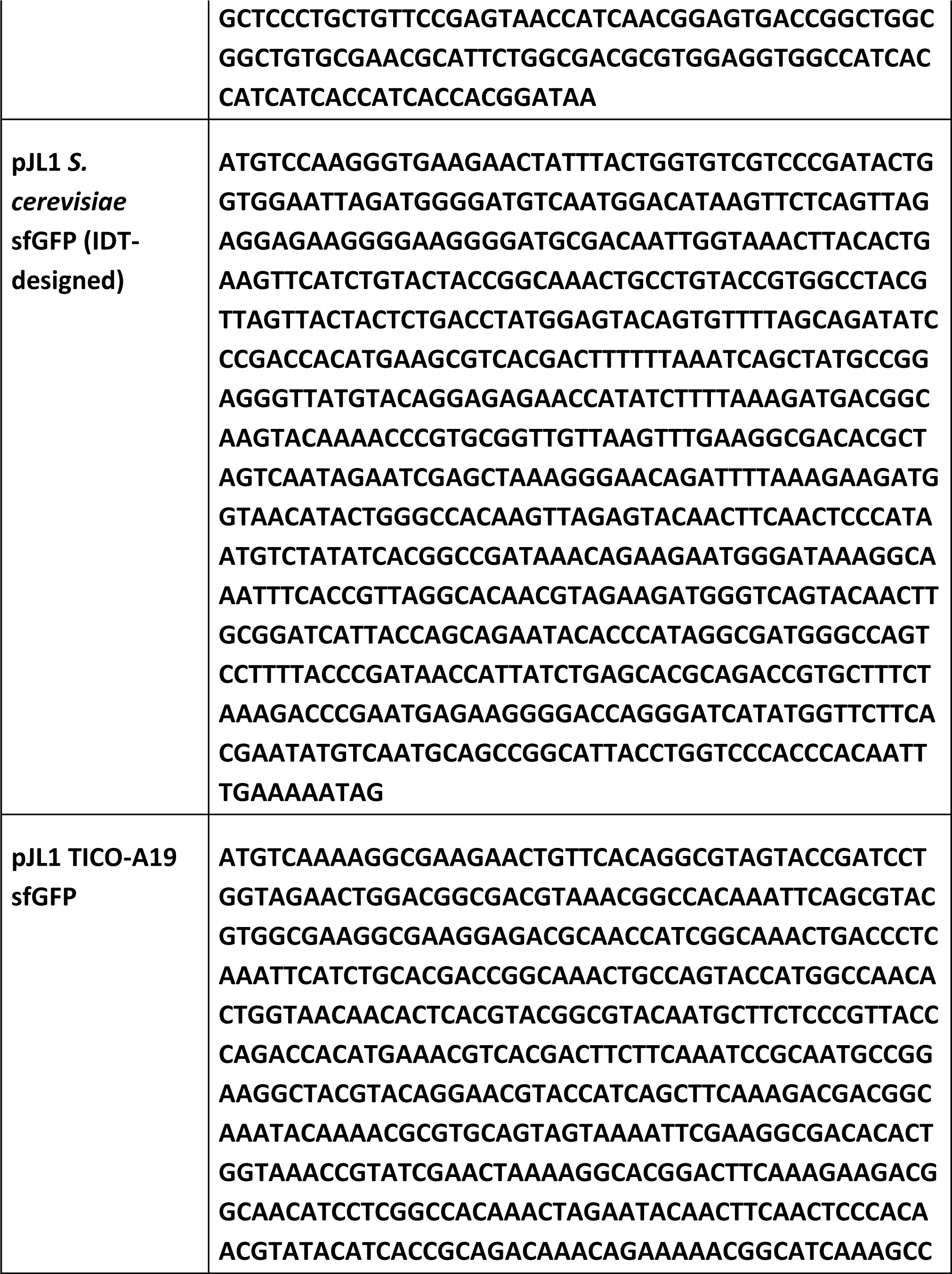
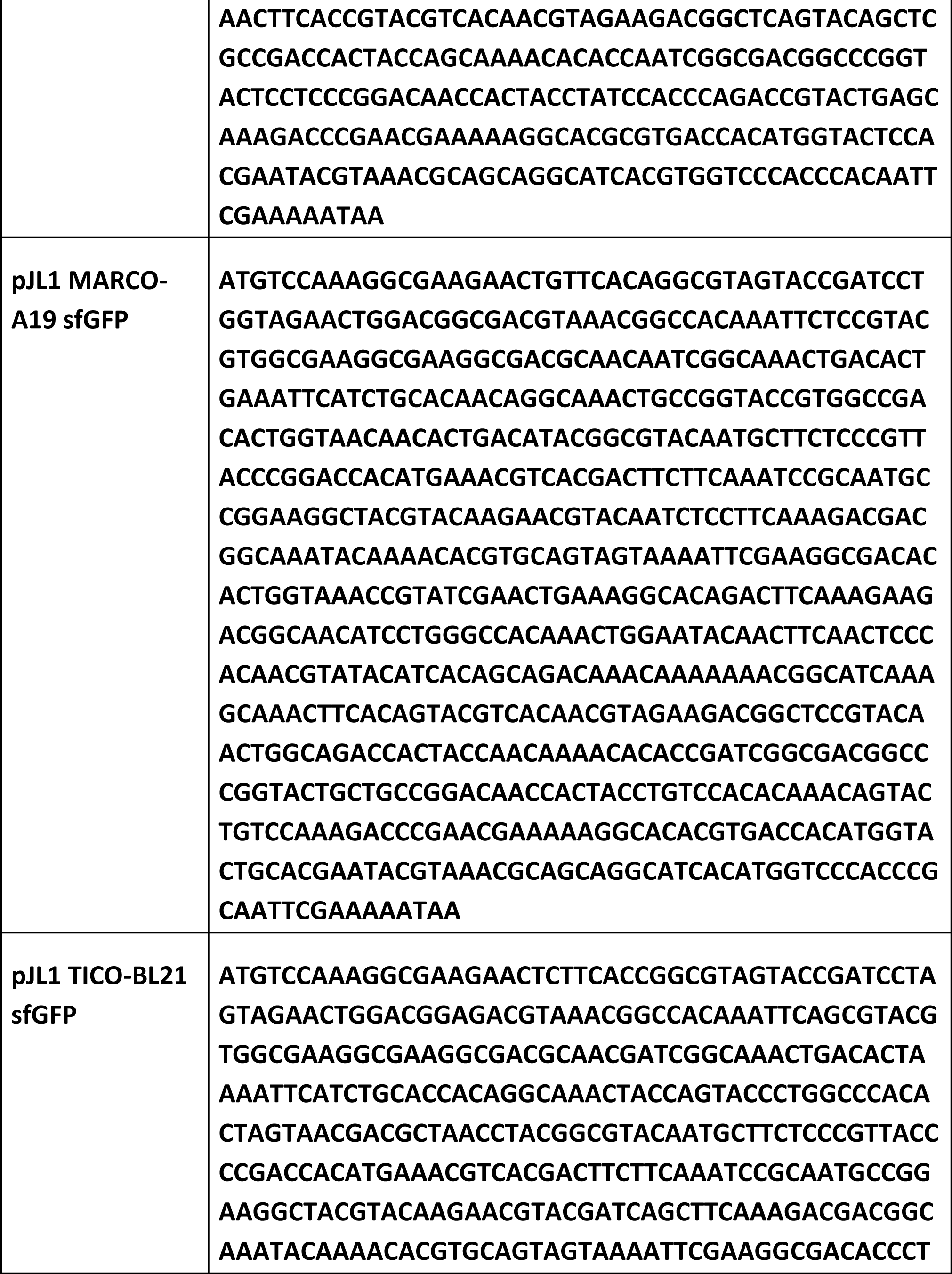
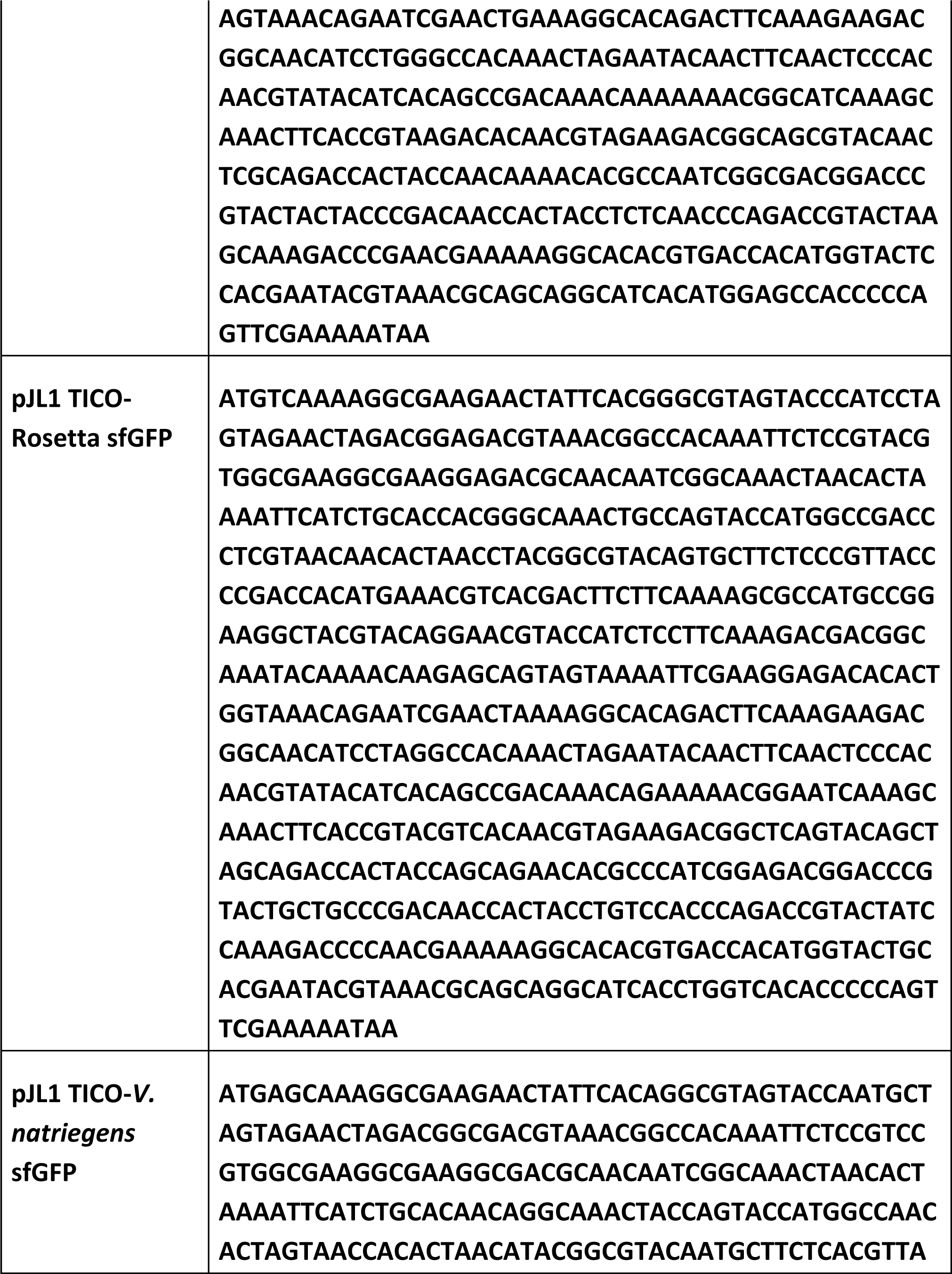
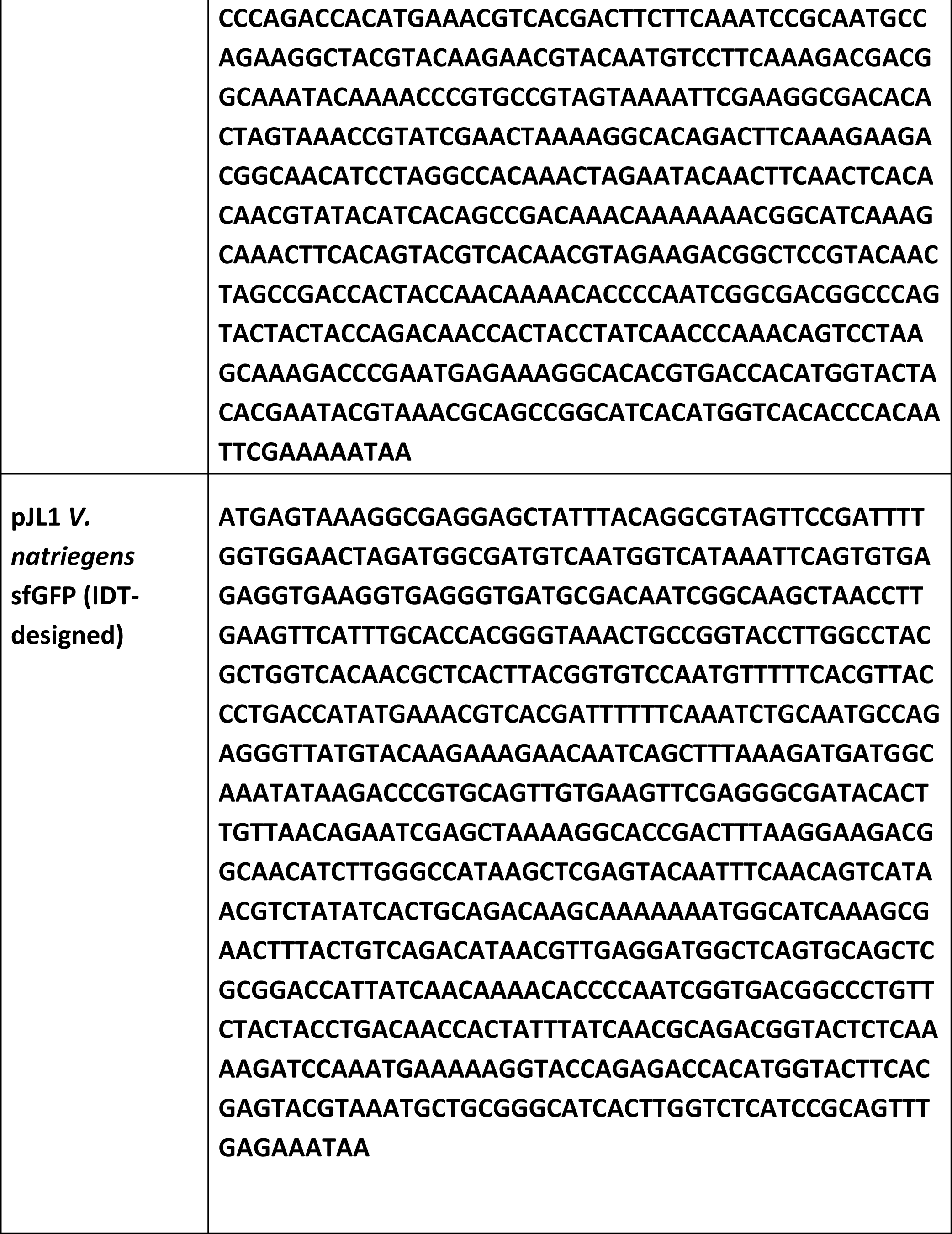
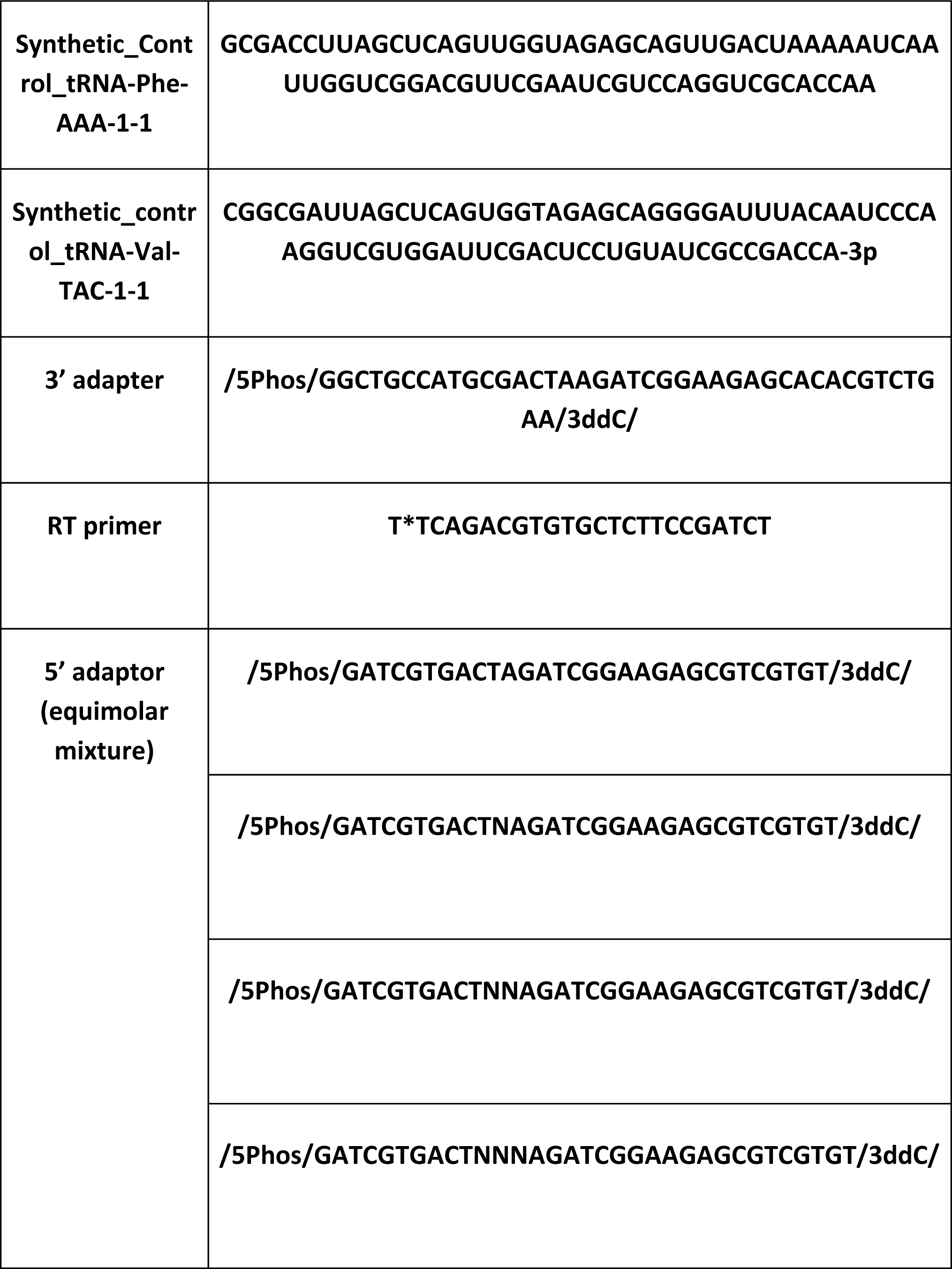
Sequences used in this study.

**Figure S1.**
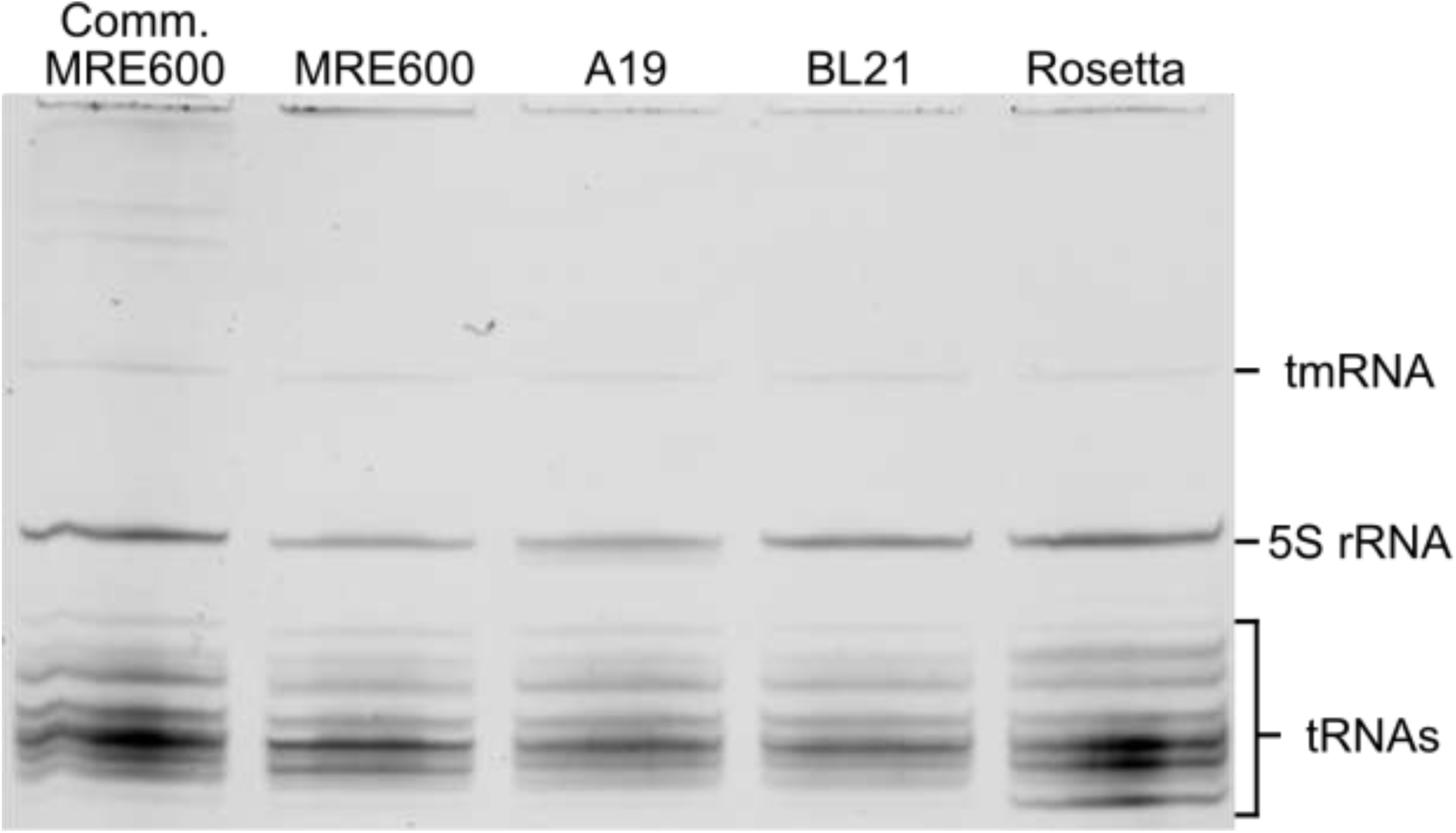
Urea-PAGE of isolated *E. coli* tRNAs. Urea-PAGE of small RNAs isolated from *E. coli* MRE600, A19, BL21, Rosetta. Urea-PAGE reveals three populations of small RNAs recovered, bulk tRNAs(70-90 nts), 5S ribosomal RNA (120nt), and transfer-messenger RNA (363 nts).

**Figure S2.**
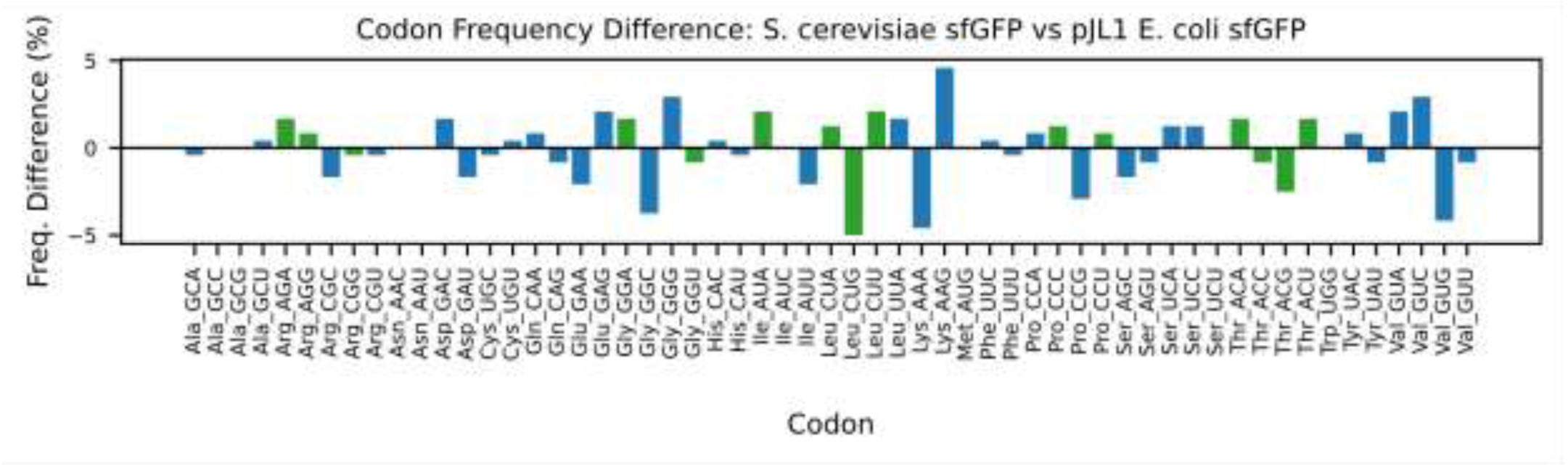
Codon frequency differences of *S. cerevisiae* sfGFP to pJL1 *E. coli* sfGFP. **Codon optimization to *S. cerevisiae* sfGFP alters codon usage frequency.** Codon frequency changes from commercially optimized pJL1 *S. cerevisiae* sfGFP relative to pJL1 *E. coli* sfGFP. Green bars illustrate codon frequency changes at codons that correspond to tRNAs provided by the pRARE2 plasmid in Rosetta.

**Figure S3.**
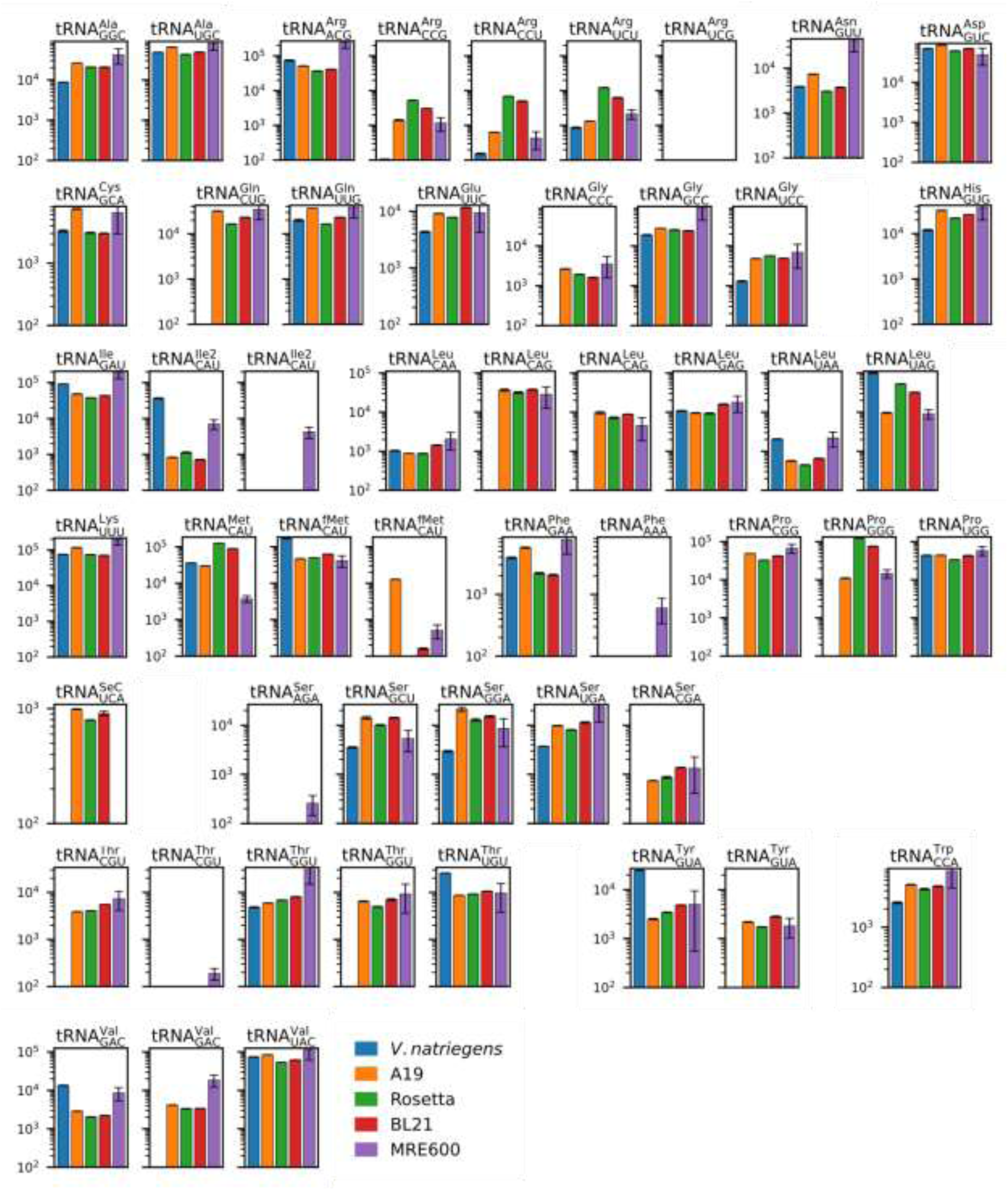
tRNA-seq reveals differences in tRNA abundances across strains. tRNA abundances of individual tRNAs given in reads per million (RPM) for each strain in this study. tRNAs are grouped within isoacceptor families. Individual tRNA reads were mapped to predicted tRNA gene sets of each strain within the study.

**Figure S4.**
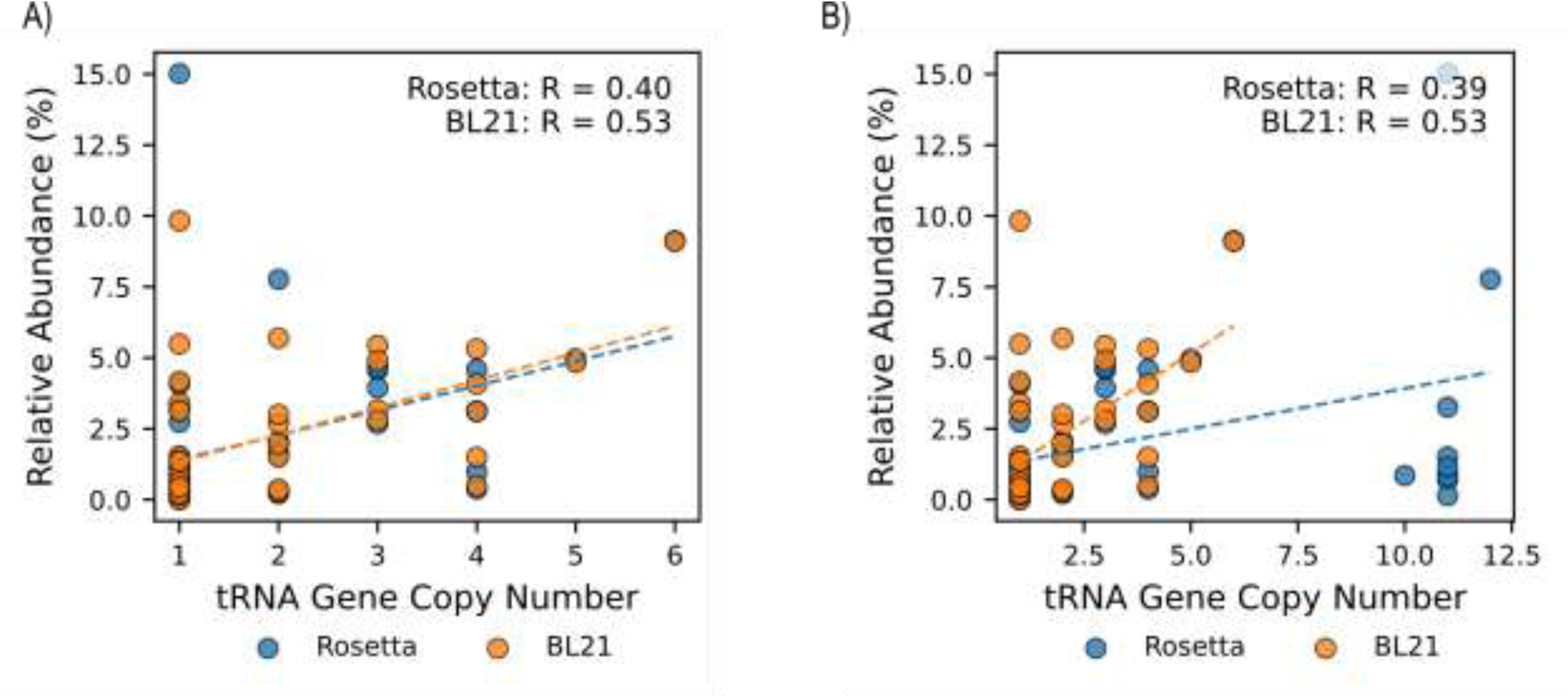
Rosetta tRNA abundance correlation plots accounting for pRARE2 gene copy number. pRARE2 gene copy fails to change correlations plots for tRNA abundance to tGCN. A) Pearson correlation plots of tRNA abundances of BL21 and Rosetta tRNA pools against the shared BL21(DE3) genome tGCN. B) Pearson correlation plots of tRNA abundances of BL21 and Rosetta tRNA pools against the shared BL21(DE3) genome tGCN with the supplementation of gene copies provided by the pRARE2 copy number.

**Figure S5.**
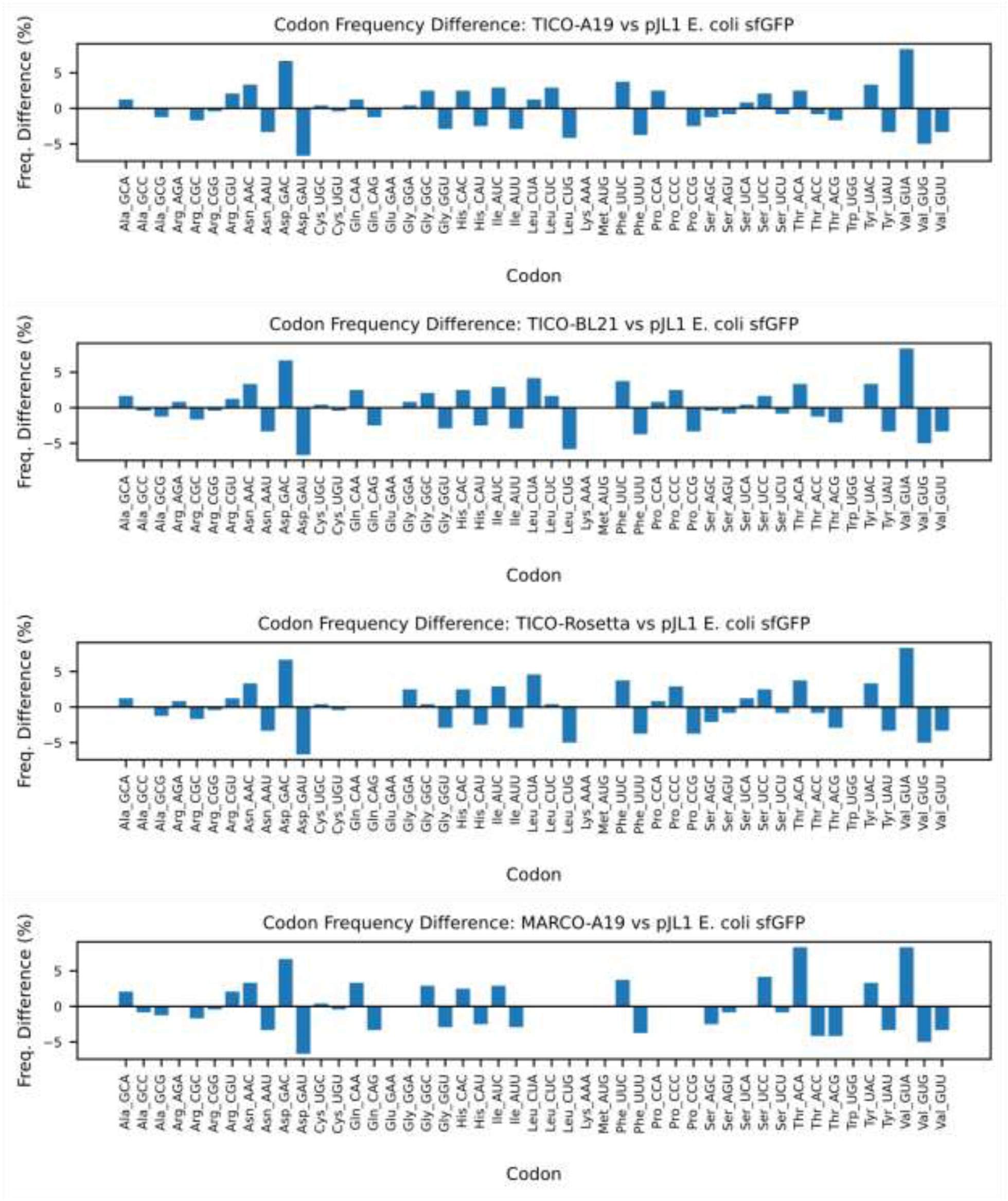
Codon frequency changes for TICO-designed and MARCO sfGFP constructs. Codon frequency changes from TICO-designed and MARCO A19 sfGFP constructs relative to pJL1 *E. coli* sfGFP. TICO optimizes coding sequences by aligning tRNA abundances of each strain to codon usage. To simplify the design, codons were chosen to be cognate Watson and Crick matches to tRNA anticodons, excluding codon selections which permit wobble decoding. This results in subtle changes of codon choices across strains, but large changes relative to pJL1 *E. coli* sfGFP. For example, Asp codons are decoded by a single tRNA^Asp^_GUC_ which matches to a cognate GAC anticodon. In pJL1 *E. coli* sfGFP, with only one exception, all Asp residues are encoded by a GAU codon. Both TICO and MARCO optimization flips codon preferences to Asp amino acids in sfGFP to be decoded exclusively by GAC codons. More subtle changes are observed within tRNA families containing diverse sets of isoacceptor abundances. For example, six Leu codons are decoded by several tRNA^Leu^, but pJL1 *E. coli* sfGFP exclusively contains CUG codons for leucine. In TICO-designed constructs, Leu codons are distributed differentially across three codons (CUA, CUC, and CUG) to engage with tRNA^Leu^_UAG_, tRNA^Leu^_GAG_, and tRNA^Leu^_CAG_ approximate to their abundances determined form tRNA-seq. In contrast, MARCO A19 construct is designed to exclusively engage with only the most abundant tRNAs within isoacceptor families. Thus, MARCO A19 mimics the exclusivity of pJL1 *E. coli* sfGFP for CUG codons to decode leucine.

**Figure S6.**
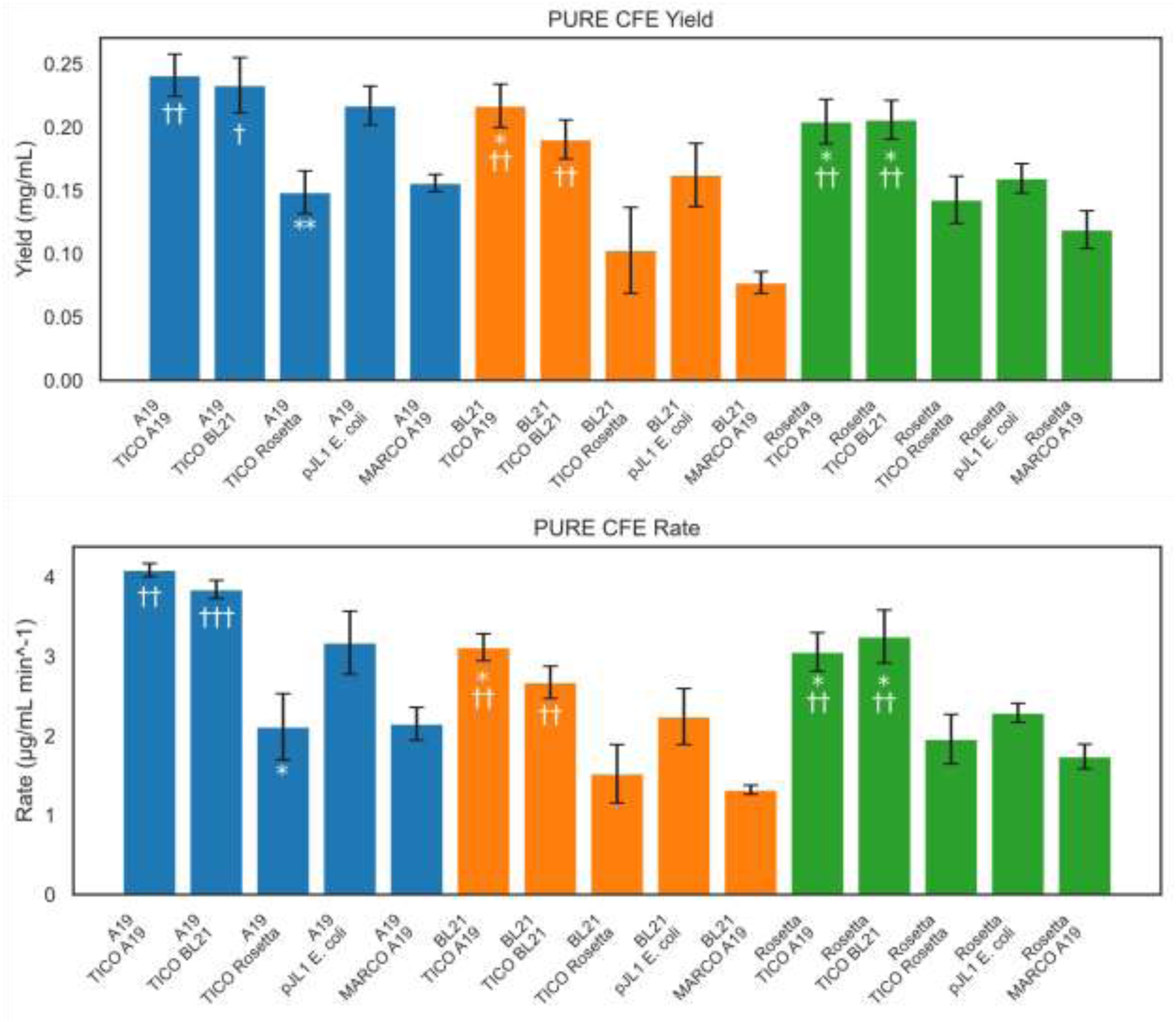
Histogram of translation yields and rates for tRNA pools translating TICO and MARCO sfGFP constructs. Translation yield and translation rates for PURE CFE reactions translating TICO and MARCO A19 sfGFP constructs using isolated A19, BL21, and Rosetta tRNA pools. Asterisks (*) represent p-values corresponding to statistical significance relative to pJL1 *E. coli* sfGFP (* = p < 0.05, ** = p < 0.01) while daggers (†) represent statistical significance against MARCO A19 sfGFP († = p < 0.05, †† = p < 0.01, ††† = p < 0.001). Error bars represent standard deviation of mean for three experimental replicates.

**Figure S7.**
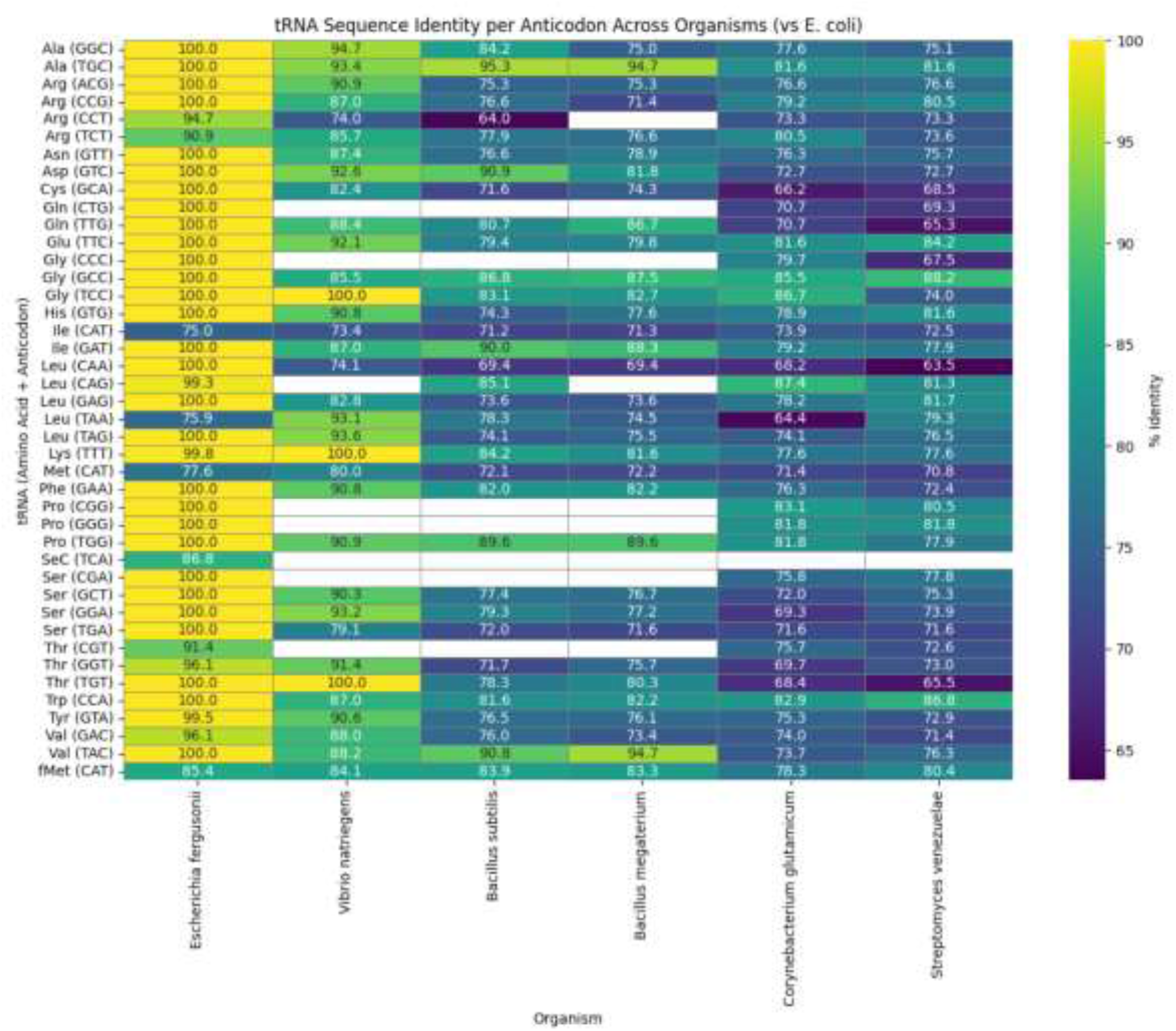
Pairwise sequence alignments of tRNA sequences of common prokaryote lysates for S30 CFE against *E. coli* K-12 MG1655 tRNAs. Pairwise sequence alignments of tRNAs from commonly used prokaryotic CFE lysates compared to *E. coli* MG1655 tRNAs pools. tRNA that are absent in these organisms but found in *E. coli* are left blank.

